# Hippocampal dentate gyrus coordinates brain-wide communication and memory updating through an inhibitory gating

**DOI:** 10.1101/2020.07.14.202218

**Authors:** José María Caramés, Elena Pérez-Montoyo, Raquel Garcia-Hernandez, Santiago Canals

**Affiliations:** Instituto de Neurociencias de Alicante (CSIC-UMH), Sant Joan d’Alacant, Alicante, Spain

## Abstract

Distinct forms of memory processing are often causally identified with specific brain regions, but a key facet of memory processing includes linking separated neuronal populations. Using cell-specific manipulations of inhibitory neuronal activity, we discovered a key role of the dentate gyrus (DG) in coordinating dispersed neuronal populations during memory formation. In whole-brain fMRI and electrophysiological experiments, we found that parvalbumin (PV) interneurons in the DG control the functional coupling of the hippocampus within a wider network of neocortical and subcortical structures including the prefrontal cortex (PFC) and the nucleus accumbens (NAc). In a novel object-location task, regulation of PV interneuron activity enhanced or prevented memory encoding and, without effect upon the total number of task activated c-Fos+ cells, revealed a correlation between activated neuronal populations in the hippocampus-PFC-NAc network. These data suggest a critical regulatory role of PV interneurons in the dentate gyrus in brain-wide polysynaptic communication channels and the association of cell assemblies across multiple brain regions.

## Introduction

Memory is thought to be encoded through modifications in the weights of synaptic connections. The circuits enabled by these synaptic modifications, and the corresponding activated cell assemblies, form brain-wide memory engrams that hold specific memory (*1*). In the dentate gyrus (DG), a region of the hippocampal formation important for pattern separation and contextual learning (*2*–*4*), activity-tagging via immediate early gene (c-Fos)-dependent expression of channelrhodopsin, has been shown to recall a contextual fear memory upon light activation (*5*). However, the same manipulation targeted to the CA1 region of the hippocampus was ineffective (*5*), despite this region being required for contextual fear memory encoding and consolidation (*6, 7*). This dissociation raises the intriguing possibility of a hitherto unknown role of the DG in memory formation, even when the expression of memory requires a systems level interaction of multiple brain regions (*8*– *10*). Key support for this idea comes from brain-wide imaging techniques which have revealed that long-term potentiation (LTP) of perforant pathway inputs to the DG not only causes measurable change in DG itself, but also enhances the functional coupling of a network of mesolimbic neocortical and subcortical structures important for memory formation (*11*–*13*). Specifically, changes in activity measured using BOLD show striking changes in frontal cortex and other regions remote from the sites of plasticity.

Studies on memory formation have largely focussed on excitatory synapses and their potentiation following repeated or coincident activation as the fundamental cellular mechanism supporting the associations between information streams (*14*–*18*). However, the contribution of inhibition has long been proposed as well (*19*), with pharmacological interventions increasing inhibitory activity associated with impaired hippocampus-dependent memory acquisition (*20, 21*) and *vice versa* during mildly decreased inhibitory activity (*22– 24*). More recent work with cell-specific manipulations and electrophysiological recordings of identified cell populations, have unambiguously demonstrated transient periods of neuronal disinhibition during learning, causally linked to memory encoding and/or expression (reviewed in (*25, 26*). Even persistent reductions of PV+ cell inhibition in DG, measured as a decrease in PV staining, has been associated with enhanced spatial learning in the watermaze (*27*). However, in the same study, a stronger PV+ inhibitory network was shown to develop in the course of memory formation as more stable memories formed (*27*). These data suggest that changes in the apparent tight coupling of excitation and inhibition (E-I) could be a fundamental network property that gates memory formation. If so, transient changes in E-I balance could lead to increased propagation of activity through regionally disbursed cell-assemblies. This would account for the brain-wide changes in BOLD seen in functional imaging.

These considerations led us to investigate the impact of selective manipulations of perisomatically innervating PV+ interneurons in the DG on memory formation and the potential brain-wide network activity that accompanies it. In combined fMRI and electrophysiological experiments, we found that pharmacogenetic control of PV-cell firing can enhance or preclude activity propagation within the hippocampus and also a brain-wide network of cortical and subcortical structures including the PFC and NAc. Disinhibition of DG enhanced the encoding of novel object-location associations whereas increased inhibition blocked it. Cell assemblies in the hippocampus-PFC-NAc network that were activated by learning were also bi-directionally controlled by DG PV cell activity. Interestingly, DG disinhibition that increased memory encoding concomitantly enhanced the functional coupling between c-Fos+ cell assemblies, but did not change the total number of activated granule cells, preserving the sparseness of activation and thus maintaining context specificity. Complementarily, the increase in PV-cell activity in the DG, decorrelated cell assemblies, without affecting their size, and prevented memory updating. Because manipulations of perisomatic inhibition were also shown to leave synaptic plasticity and dendritic integration unaltered in granule cells, our results suggest that PV-cells in DG are a major contributor of a system-level gating mechanism that orchestrates neuronal activity in separated brain regions.

## Results

### DG PV cells regulate granule cell output bidirectionally

We regulated PV-cell activity in the DG with Designer Receptors Exclusively Activated by Designer Drugs (DREADDs) (*28*). These receptors allow gain- or loss-of-function manipulations of PV-cell firing, depending on the subtype of receptor expressed, without imposing an external pattern of activation nor completely blocking spiking. We injected stereotaxically adeno-associated virus 5 (AAV5)-human synapsin1 (hSyn)-DIO-hM4D(Gi)-mCherry in the hilus of the DG of PV-cre transgenic mice (129P2-Pvalb^tm1(cre)Arbr^/J) to express the inhibitory hM4D(Gi) receptor in dentate PV interneurons (Fig. 1A; referred from now as PV-Gi animals). Conversely, for the activation of PV-cells we injected the AAV5-hSyn-DIO-hM3D(Gq)-mCherry virus to express the excitatory hM3D(Gq) receptor (Fig. 1A; referred as PV-Gq animals). The proportion of PV+ interneurons infected in the DG was 92.8 ± 3.8 % and 91.2 ± 3.5 % for hM4D(Gi) or hM3D(Gq) viruses, respectively (Supp. Fig. 1A-C).

**Fig. 1.**
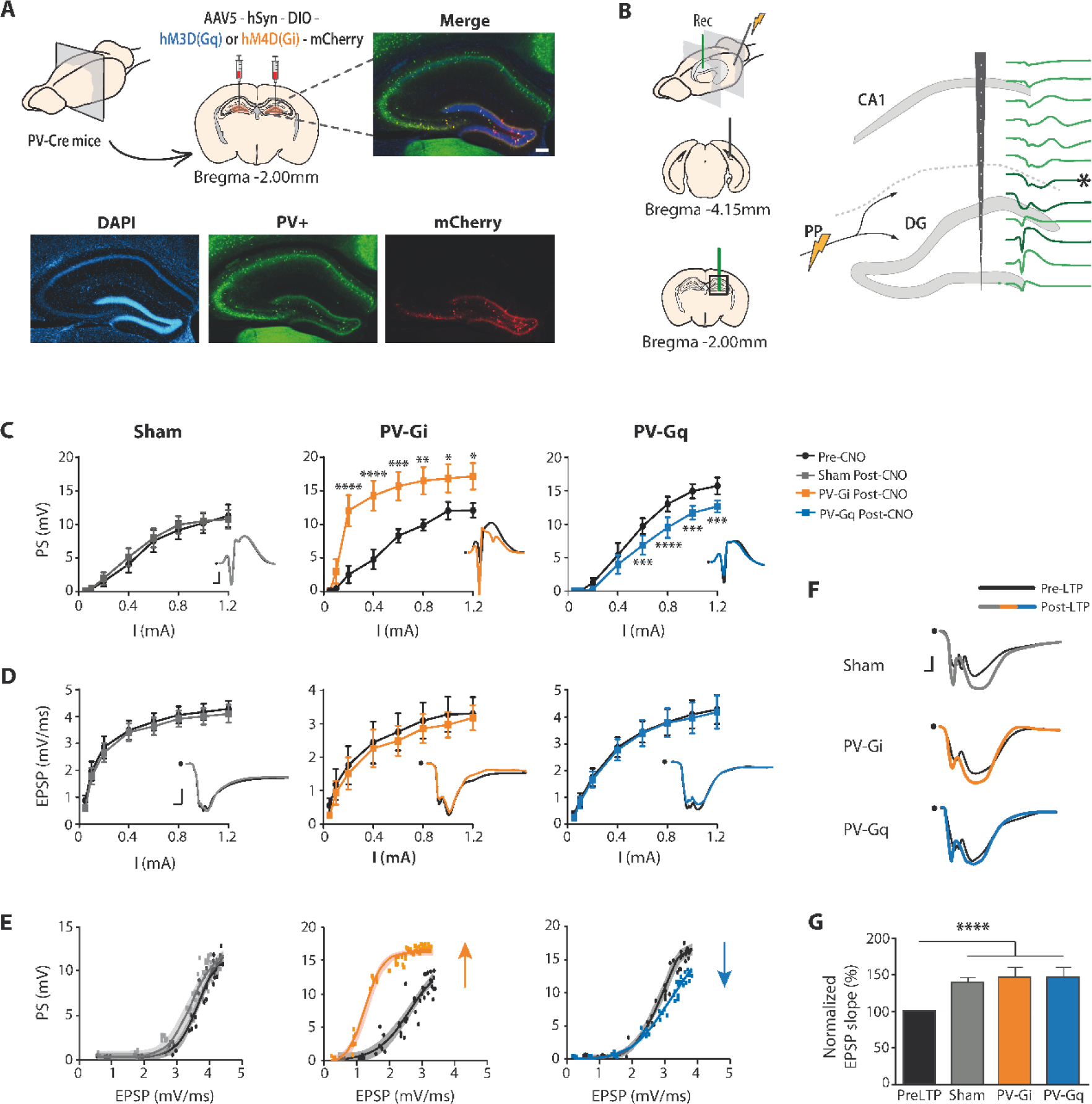
Modulation of PV-cell activity in the DG controls GC output preserving synaptic activity and plasticity. **A**, DG virus injection and double labelling of PV+ interneurons (taken from a PV-Gq animal). Scalebar = 200µm. **B**, Multichannel *in vivo* electrophysiological recordings (green traces) in response to perforant path (PP) stimulation; Thick traces indicate the selected channels for PS analysis (lower trace) and EPSP (upper two traces). For PV-Gi animals, the larger and faster PS interfered with EPSP measures in the optimal location and a more distal recording is taken (asterisk, see Material and Methods section). **C**, Comparison of PP stimulus-response curves of DG PS amplitude before (black) and after CNO i.p. injection in Sham (grey), PV-Gi (yellow) and PV-Gq (blue) animals. Insets: representative PS waveforms (scale 2 ms, 4 mV). **D**, Comparison of PP stimulus-response curves of DG EPSP slope before and after CNO i.p. injection (same colour-code as before). Insets: representative EPSP waveforms (scale 1 ms, 1 mV). **E**, Input-Output (EPSP vs. PS) curve demonstrating increased (PV-Gi) and decreased (PV-Gq) granule cell output for equal synaptic inputs (same colour-code; non-linear regression fit, shadow represents 95% CI). **F and G**, LTP induction. After CNO injection, LTP induction enhanced the EPSP in all groups (**F**). Quantitative analysis showed no differences between groups (**G**). Pre-LTP data are represented together for simplicity and EPSP values normalised vs. pre-LTP levels. Group data represent mean ± SEM. All statistical values are detailed in Supp. Table 1. *p≤0.05, **p≤0.01, ***p≤0.001, ****p≤0.0001.

In PV-Gi animals *in vivo*, evoked potentials in the DG following stimulation of the perforant pathway, the main entorhinal cortex (EC) input to the hippocampus (Fig. 1B), showed a significant increase in the amplitude of the population spike (PS) after CNO (i.p. 1 mg/Kg) administration (Fig. 1C), reflecting the facilitated firing of GC. Conversely, decreased PS amplitude in the DG (Fig. 1C) was recorded in PV-Gq animals upon CNO administration, indicative of the hindered GC firing. These effects were maximal 30 min after CNO administration and remained constant for the 3 hr of the remainder of recording (Supp. Fig. 1D) (*28*). Importantly, the excitatory postsynaptic potentials (EPSPs), reflecting the synaptic responses and dendritic integration in GC, were unchanged in all groups (Fig. 1D). This was clearly evidenced in the input-output curves relating synaptic inputs to spike output (Fig. 1E), demonstrating the selective control of PV-cells over the GC output. The integrity of the dendritic function was further demonstrated in synaptic plasticity experiments in which the induction of long-term potentiation (LTP) by high-frequency stimulation of the perforant pathway was unaffected by CNO administration in any of the experimental groups (Fig. 1F and G and Supp. Fig. 1E).

### DG PV cells bidirectionally control memory updating

We then investigated whether the gating of GC’s output, keeping an intact synaptic and dendritic function, had any impact on memory formation. We used the hippocampal-dependent novel object location task (NOL) (*29, 30*). In this two-phase task, mice first learn new spatial cues (2 identical objects) in an otherwise familiar context during a single 10 min exploration trial. They are then allowed to retrieve the encoded memory of object locations 24 h later in the same familiar context with 1 of the 2 objects displaced. If memory is intact, this elicits a relative exploratory preference towards the moved object (See Methods and Supp. Fig. 2A). Activity of PV cells in the DG was modulated as before (i.p. 1 mg/Kg CNO) either 90 min before the exploration trial (encoding phase), 10 min after the exploration trial when the animal is returned to its home-cage (consolidation phase) or 90 min before the retrieval test 24 h after encoding (retrieval phase) (Fig. 2A). The results demonstrate a highly significant and bidirectional effect of modulating PV-inhibition in the DG during memory encoding (Fig. 2B), but not during consolidation (Fig. 2C) or retrieval (Fig. 2D). Decreasing perisomatic inhibition during encoding resulted in enhanced performance 24 h later, while increasing the inhibitory tone prevented encoding (Fig. 2B). Locomotor and anxiety measures during field exploration and the elevated plus maze, respectively, were indistinguishable between groups (Supp. Fig. 2C and E). An important control was to repeat the NOL task in the same group of animals and substitute CNO administration by its vehicle (saline) in the encoding phase; comparable memory encoding was observed in all groups (Fig. 2E).

**Figure 2.**
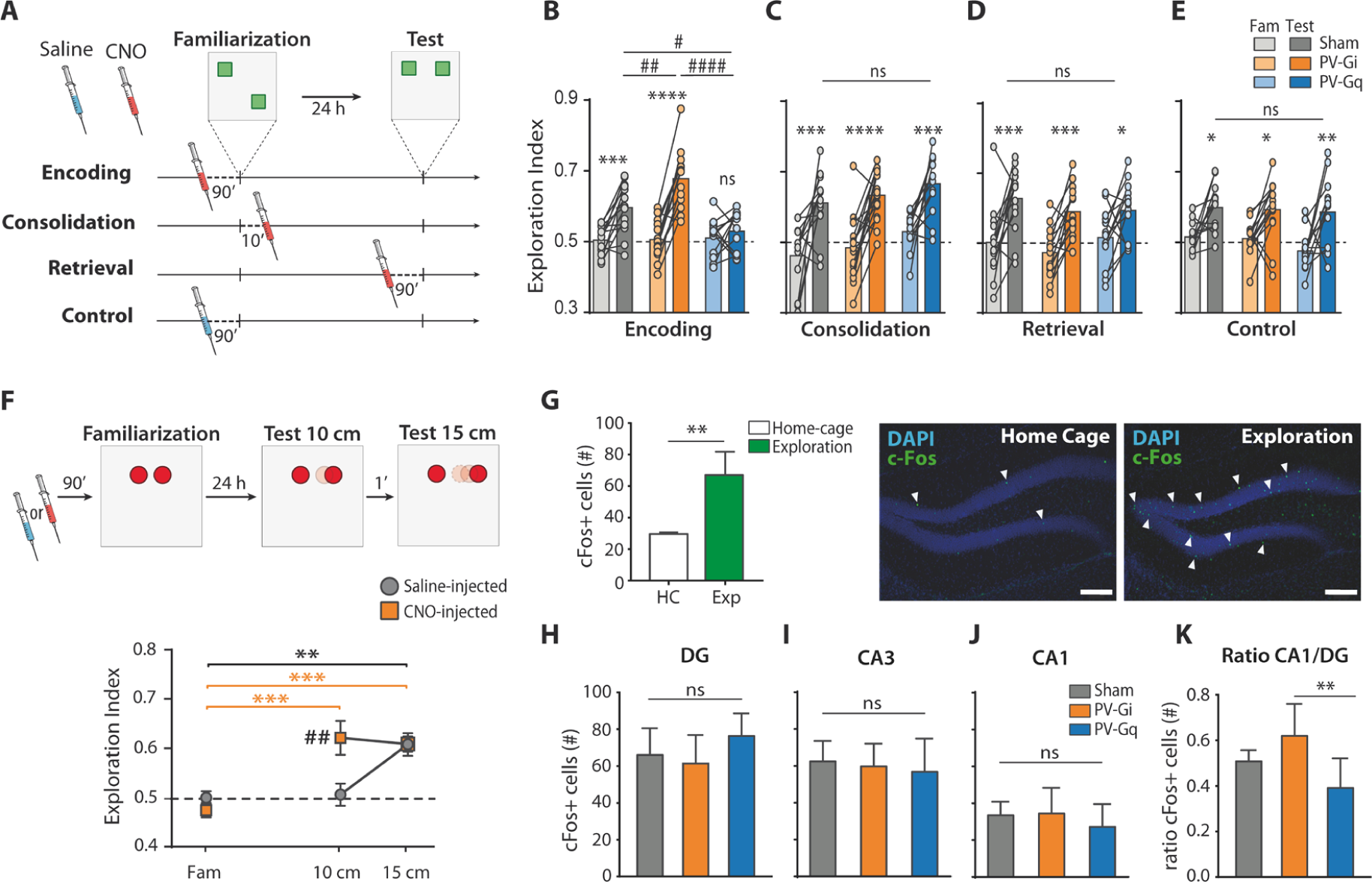
Memory encoding under the control of DG PV-cells. **A**, NOL protocols used to modulate PV-cell activity in different task phases. For the control experiment, the same animals were saline-injected (no PV-modulation). **B to E**, Performance in the NOL task with interventions in encoding (**B**), consolidation (**C**) or retrieval (**D**), and the saline control (**E**). Values higher than 0.5 denote preference towards the moved object. Dots represent pair observations of individual animals and bars mean values. Light and dark colours refer to familiarization and test, respectively. * = within group (Fam. vs. test), # = between group (Sham vs. PV-Gi vs. PV-Gq). **F**, Enhanced novelty detection. The object is first displaced only 10 cm, and then 5 cm more to the final position at 15 cm from the initial location (top). CNO (yellow) or saline (grey) is injected in the familiarization phase. The same animals were used for both conditions (bottom). Data represent mean ± SEM. * = within group comparison (Fam. vs. test), # = CNO vs. saline. **G**, Activated c-Fos+ cells in the DG during the NOL familiarization phase vs. home-cage (HC). Insets: c-Fos immunostained cells (white arrows) in the DG (right). Histograms represent mean ± SEM. Scale bars: 100 µm. **H to J** Number of c-Fos+ cells in Sham (grey), PV-Gi (yellow) and PV-Gq (blue) animals, in the DG (**H**), CA3 (**I**) and CA1 (**J**), respectively. **K**, Ratio of c-Fos+ cells activated in CA1 as a function of DG activation (CA1/DG) in each subject and averaged by group (mean ± SEM). * = PV-Gi vs. PV-Gq. All statistical values are detailed in Supp. Table 2. *#p≤0.05, **##p≤0.01, ***p≤0.001, ****####p≤0.0001.

One hypothesized role for the DG in memory formation is its contribution to pattern separation (*2*–*4*), a function that would theoretically benefit from the known sparse activation of GCs (*2, 3*). Compatible with this role, enhanced object location discrimination was observed in PV-Gi animals injected with CNO as the difficulty of detecting the magnitude of object displacement was made harder (Fig. 2F). A small environmental change not ordinarily detected in control conditions was noticed and effectively assimilated into memory when PV-cells were inhibited. However, previous work manipulating the activity of somatostatin positive interneurons in the DG showed an enlargement of the recruited GC assembly (*31*), which would work against the sparsity of GC activation and their capacity to discriminate patterns. To clarify this issue, we used cellular imaging of activity-dependent c-Fos expression, and asked whether specific PV-interneuron activation or inactivation had an impact on the number of cells recruited in the local assembly. The baseline sensitivity of this assay is shown by the number of c-Fos+ cells in the DG being increased by object exploration in the NOL task compared to those of home-cage mice (Fig. 2G). The logically appropriate comparisons are between the number of c-Fos+ GCs in CNO injected PV-Gi, PV-Gq animals and CNO injected PV-Sham controls that revealed a constant size of the recruited GC assembly across conditions (Fig. 2H). As a control, in the same animals, we validated the successful DREADDs manipulation showing enhanced activation of PV+ interneurons in PV-Gq animals, and *vice versa* in PV-Gi (Supp. Fig. 3). Thus, although GC spiking probability is radically changed, the number of c-Fos+ cells in the DG seems to be regulated by the synaptic input, unaltered here, rather than overt neuronal firing. In addition, neither the number of c-Fos+ pyramidal neurons in CA3 (Fig. 2I) nor CA1 (Fig. 2J) were altered by PV-cell modulation in the DG. Therefore, the sparseness of the hippocampal cell assemblies was preserved regardless of the PV-inhibitory level in the DG and the number of c-Fos+ cells thus dissociated from memory performance.

**Figure 3.**
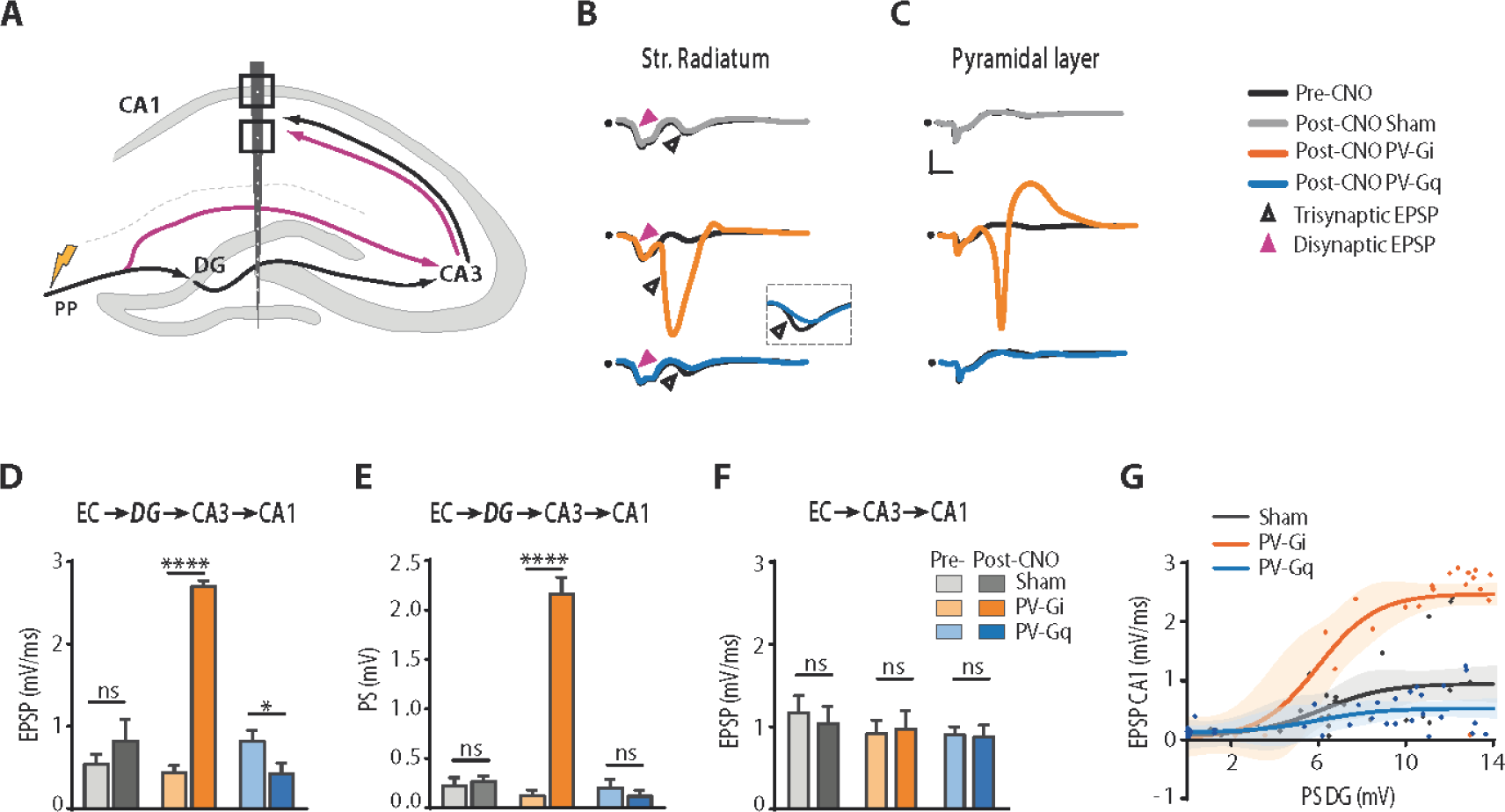
CA1 output is strongly regulated by DG PV-cell activity. **A**, Schematic representation of the trisynaptic (EC→DG→CA3→CA1; black) and disynaptic (EC→CA3→CA1; purple) hippocampal circuits. Squares mark the position of the recordings in CA1 stratum radiatum (lower) and pyramidale (upper). **B, C**, Evoked potentials in CA1 in response to PP stimulation showing trisynaptic and disynaptic EPSPs in stratum radiatum (**B**) and the trisynaptically evoked PS in the pyramidal layer (**C**). **D to F**, Quantitative group analysis of the trisynaptic EPSPs (**D**) and PS (**E**) and the disynaptic EPSP, before (light colours) and after (dark colours) CNO injection. Data represent mean ± SEM. **G**, Activity transfer from the DG to CA1. DG PS amplitude is plotted against their trisynaptically evoked EPSPs in CA1 (solid lines represents non-linear regression fit, shadow represents 95% CI). All statistical values are detailed in Supp. Table 3. *p≤0.05, ****p≤0.0001.

Importantly, however, the inter-subject c-Fos expression variability analysis indicated that the functional coupling between the c-Fos+ populations inside the hippocampus was bidirectionally modulated by DG inhibition (Fig. 2K and see Methods). The number of c-Fos+ neurons in CA1 and the DG largely covaried in all groups (Pearson correlation coefficients of 0.94, 0.92 and 0.88, for sham, PV-Gi and PV-Gq, respectively), but the number of c-Fos+ cells recruited in CA1 per active DG cFos+ cell was increased by perisomatic disinhibition, consistent with enhanced activity propagation from DG to CA1 and in parallel with enhanced memory encoding. In contrast, increased perisomatic inhibition was sufficient to decrease this ratio and prevent memory formation.

### CA1 output is strongly regulated by DG PV cells

We then used *in vivo* multi-site electrophysiological recordings to investigate intra-hippocampal functional connectivity (Fig. 3A). In control conditions, stimulation of the perforant pathway with a single electrical pulse produced the well-known activity propagation from DG to CA1 in the trisynaptic circuit (EC→DG→CA3→CA1), generating only small synaptic currents in the *stratum radiatum* (Fig. 3C and D) that were alone insufficient to drive action potential firing in CA1 pyramidal neurons (Fig. 3C and E) (*32*). However, during PV cell inhibition in the DG, the same stimulation triggered strong synaptic responses and robust firing (Fig. 3B to E). Increasing the activity of PV-cells produced the opposite effect (Fig. 3B to E). The relevant control here was to quantify separately the disynaptic propagation from CA3 to CA1 (EC→CA3→CA1), bypassing the DG, to check on the absence of differences within or between groups before and after CNO administration (Fig. 3B and F and Supp. Fig. 4).

**Fig. 4.**
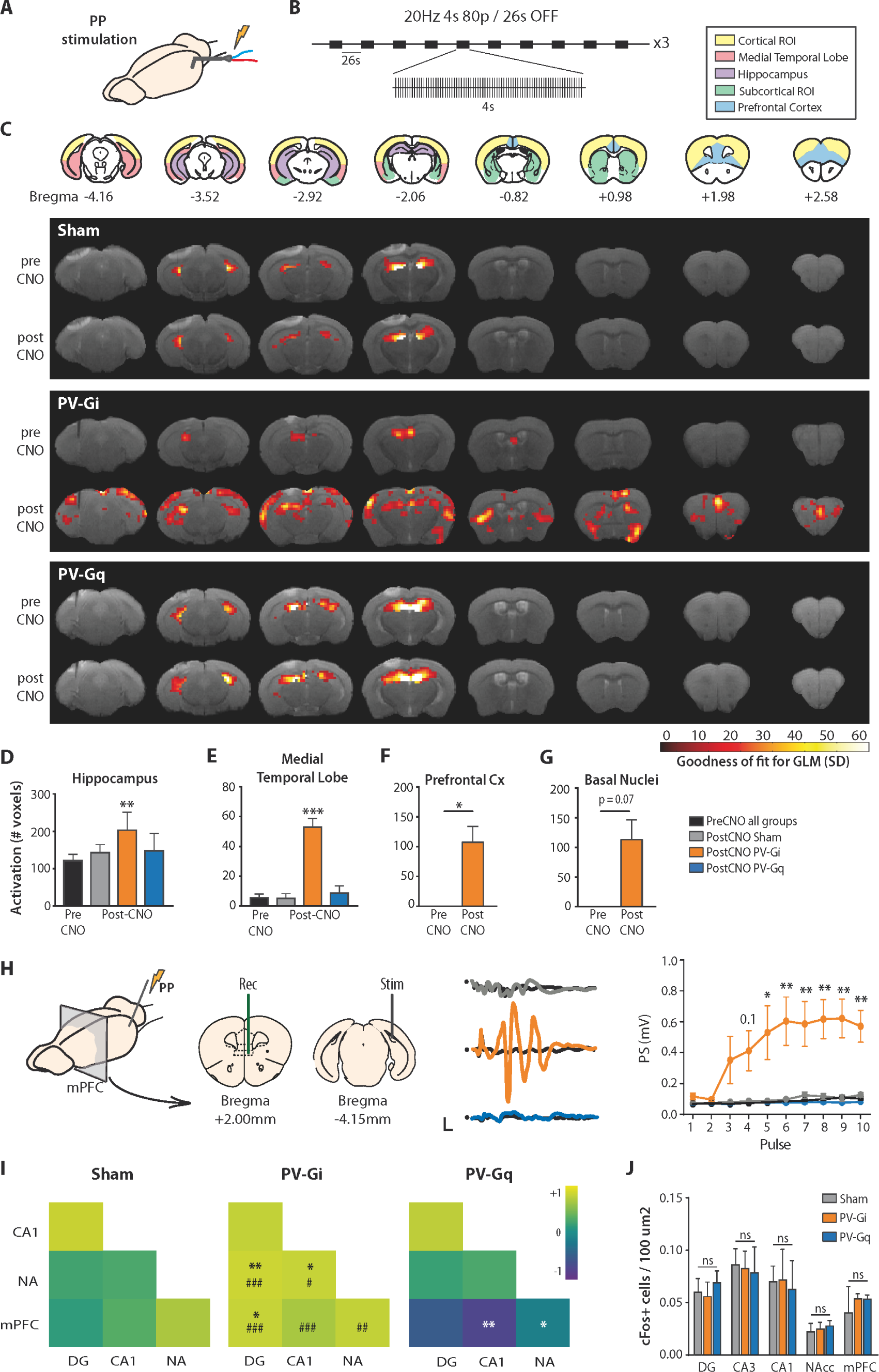
DG PV-cells gate long-range communication channels from the hippocampus. **A**, Schematic of the preparation. **B**, 20 Hz stimulation protocol (80 pulses per train) applied in the PP. **C**, Upper: regions of Interest (ROIs) used in the quantitative analysis. Numbers indicate mm from Bregma. Lower: functional maps evoked by the stimulation before and after CNO injection, and overlaid on T2 anatomical images. Colour-code represents the goodness of fit of the GLM analyses thresholded at p<0.01. **D to G**, Number (mean ± SEM) of statistically significant active voxels per group and condition in the indicated ROIs. Pre-CNO data are represented together for simplicity. **H**, Evoked electrophysiological potentials in the PFC in response to PP stimulation. Stimuli consist in 1 Hz trains of 10 pulses. Scale bars: 5 ms, 0.1 mV. Group data represent mean ± SEM. Pre-CNO responses for the three groups are averaged for simplicity (black). **I**, Co-variation of c-Fos+ expression in the hippocampus, PFC and NAc. Colour-code refers to Person-correlation coefficients. * = Sham vs. PV-Gi or PV-Gq; # = PV-Gi vs. PV-Gq. **J**, Number (mean ± SEM) of c-Fos+ cells per 100 µm^2^ activated during the NOL familiarization phase in the indicated ROIs. All statistical values are detailed in Supp. Table 4. *#p≤0.05, **##p≤0.01, ***###p≤0.001.

These results reveal that facilitated firing in CA1 is causally influenced by specific disinhibition of the DG relay. However, it was not only the consequence of increased GC firing, since the amplitude of the DG PS did not fully explain CA1 activity (Fig. 3G). We selected stimulation trials with comparable activation of GCs (same rage of PS amplitudes) across all experimental groups and quantified the corresponding CA1 propagation, and found that the effective activity transfer from DG to CA1 was largely increased in CNO injected PV-Gi animals (Fig. 3G). What might be the basis of this additional effect? In these experiments, it was observed that the firing delay of GCs was decreased (Supp. Fig. 5B and C) and its synchrony increased (Supp. Fig. 5D and E). Inclusion of synchrony as co-variable significantly explained the evoked CA1 EPSP (F(1,80)=4.13, *p*= 0.046), indicating a contribution to the propagation to CA1 and a role of the precise timing of GC firing.

### DG PV-cells gate long-range communication channels in the brain

The change in CA1 firing after a cell-specific alteration of DG inhibition potentially has major implications because activity in hippocampal long-range circuits largely relies on CA1 output. What then might be the systems-level consequences of the bi-directional control of DG physiology by PV-cells? A series of brain-wide functional magnetic resonance imaging (fMRI) experiments in control, PV-Gi and PV-Gq animals was performed (Fig. 4). In control animals, stimulation of the perforant pathway (Fig. 4A and B and Supp. Fig. 6A and B) activated structures of the hippocampal formation including the DG, CA3, CA1 and subiculum, with little or no extra-hippocampal propagation (*12, 33*), and with no effect of CNO (1 mg/Kg i.p.) (Fig. 4C to G). However, inhibition of PV interneurons in PV-Gi animals given CNO allowed hippocampal activity to propagate to cortical areas in the medial temporal lobe (including the entorhinal and perirhinal cortices), the PFC (including the prelimbic, infralimbic, orbitofrontal and cingulate cortices) and the retrosplenial cortex (Fig. 4C and F). Hippocampal activity also reached subcortically to the striatum, notably the NAc (Fig. 4C and G). Conversely, increasing PV-inhibitory tone in PV-Gq mice resulted in fMRI activation maps with no extra-hippocampal propagation (Fig. 4C, E to G and Supp. Fig. 6).

To validate the fMRI findings, we performed simultaneous electrophysiological recordings in the medial PFC (Fig. 4H), one of the fMRI-identified extra-hippocampal structures activated during DG disinhibition. Electrophysiological potentials evoked by stimulation of the perforant pathway were only seen when the activity of the PV-cells was decreased (Fig. 4H). Therefore, downregulating the perisomatic inhibitory tone in the DG enhances the effective connectivity from the hippocampus to the PFC. To extend this finding to additional brain regions and, importantly, under anesthesia-free conditions, we measured the covariation of c-Fos expression between the hippocampus, PFC and NAc in animals performing in the NOL task. Activation in these brain regions has been shown to be necessary during memory encoding in spatial memory tasks (*34*–*37*) and we demonstrated that encoding in the NOL task indeed activates the three brain regions in comparison to homecage exploration (Fig. 2G and Supp. Fig. 7). The important analysis here was the covariation between c-Fos+ cell assemblies, demonstrating enhanced coupling between the hippocampus, PFC and NAc in the PV-Gi animals and decreased coupling in the PV-Gq animals (Fig. 4I). Interestingly, the number of C-Fos+ cells in the NOL task across conditions and regions was constant (Fig. 4J), highlighting the effect of the experimental manipulation on the coordination between the activated cell assemblies, rather than their activation *per se*. Overall, DG PV-cells regulate the functional coupling of the hippocampus within a network of higher-order associational cortices and cortical and subcortical limbic structures known to subserve memory formation (*38, 39*).

## Discussion

The key new finding is the identification of a systems-level gating mechanism in the DG and operated by PV-interneurons, permitting effective hippocampal coordination of cell assemblies in a distributed brain network. DG disinhibition increased functional connectivity between neuronal populations in the hippocampus, associative and prefrontal cortical areas and NAc and enhanced the encoding of novel object-context associations and context discrimination, while the opposite was found when PV-cell activity was increased. We support these findings in whole-brain fMRI data and targeted electrophysiological recordings in anesthetized animals, as well as cognitive evaluation and c-Fos network analysis in awake and freely moving animals. Because the brain receives a continuous bombarding of sensory information, we propose that the capacity of the DG to functionally couple or decouple a large network of brain regions relevant for learning (*8*–*10*), might provide the flexible mechanisms required to select information and assimilate it into memory, or discard it, preserving the integrity of the memory base.

### Mesoscopic and macroscopic consequences of DG disinhibition

Our results link the effect of the E-I balance at the cellular level in the DG circuit (GC-PV interactions) to its mesoscopic (intrahippocampal) and macroscopic (brain network) levels. A first characteristic is the regulation of GC neuronal firing without affecting dendritic currents and synaptic plasticity (Fig. 1C-G), as expected form the perisomatic innervation of GCs by PV+ interneurons (*40, 41*). In addition, GCs firing became more synchronous (less jitter) and less delayed with respect to the synaptic input (Supp. Fig. 5B-E) during PV-cell inhibition. This effect might have a large impact on intrahippocampal effective connectivity, since it has been shown that the relative timing of the CA3 and EC inputs onto the apical dendrite of the CA1 pyramidal neurons determines the pyramidal cell firing probability (*42, 43*). Since activity and timing in the di-synaptic pathway to CA1 (EC→CA3→CA1) in our experiments were shown to be unaltered (Fig. 3B and 3F), as it was likely the case in the direct temporoammonic pathway (EC→CA1) far from the DG manipulation, the change in relative timing in these two pathways relative to the tri-synaptic pathway (EC→DG→CA3→CA1) during DG disinhibition, may constitute the substrate for the large increase in the CA1 output. Accordingly, both, increased firing probability and timing precision of GCs contributed to the large increase in CA1 firing during DG disinhibition, as demonstrated by the general lineal model analysis.

At the macro-scale level, enhanced CA1 output during DG disinhibition reached a network of extrahippocampal structures. This result might be partially explained by the enhanced firing of CA1 neurons *per se* (*44*), however, the increase in the correlation between c-Fos+ assemblies in the behavioural experiment, while keeping a constant number of task activated c-Fos+ cells per region, suggests a further contribution of network synchronization. It is known that PV+ interneurons are fundamental network elements generating gamma oscillations (*45, 46*) and organizing brain rhythms (*47*). Communication between brain regions is thought to occur when the oscillatory activity in connected populations is coherent or phase locked (*48*) and it has been shown that PV-cell inhibition is able to induce phase resetting (*49*). Therefore, we hypothesize that the activation level of PV-cells in the DG sets the phase of ongoing oscillations facilitating or precluding subsequent information exchange in the network. Supporting this hypothesis, a recent electrophysiological analysis of hippocampal gamma and theta activity in CA1 and DG during novelty exploration and memory guided behaviour, shows increased theta synchronization between regions associated to theta-gamma interactions (*50*). In this study, gamma activity, representing E-I circuit interaction, was associated to theta-phase shifts and synchronization between regions (*50*). Therefore, in addition to the increased gain of the CA1 output, we speculate that long-range activity synchronisation contributes to the enhanced functional coupling during DG disinhibition. This might be a special role of PV circuits in the DG, since inhibition of PV-cells in CA1 or the medial PFC disrupts the timing of hippocampal ripples and cortical spindles, respectively, decreasing their temporal coincidence and impairing memory consolidation (*7*). Overall, we have shown that a relatively simple and constrained manipulation in the local DG circuit has a large impact on the macro-scale organization of functional connectivity, supporting the view of the DG as a critical node in the brain network for memory formation (*12, 13*).

### Binding cell assemblies, updating the memory

Learning new context associations activates cell assemblies in the DG, CA3, CA1, PFC and NAc (Fig. 2G-H, Fig. 3I-J and Supp. Fig. 7). However, this activation was not sufficient for memory formation. Equally sized but functionally unbound (uncorrelated) assemblies, found during increased DG inhibition, were associated with memory failure. In contrast, enhanced functional binding between comparable cell assemblies, obtained in our experiments during DG disinhibition, was associated with improved memory outcomes. This result indicates that experience-activated cell assemblies need to be integrated into systems-level circuits to encode the memory, and point to the DG as a critical node in this network function. A second function of this circuit mechanism can be found in the preservation of stored memories from continuous overwriting, in this case by decorrelating or decoupling brain assemblies. The threshold for memory updating would thus be set by the instantaneous E-I balance in the DG, a prediction that should be tested in future experiments. The dynamic control of PV-cell activity levels in the DG appears as a critical gate for memory updating.

Our results circumscribe this DG function to the earliest stage of memory formation, at the time of initial stimulus encoding, with no detectable contribution during memory consolidation or retrieval (Fig. 2). This result is compatible with a role of the PV cells of the DG in the coordination of the initial associations between dispersed cell assemblies necessary to build up the circuits supporting a stable memory engram, likely providing the scaffold for subsequent stabilization through the consolidation process (*9, 10, 34, 51*). This could be the mechanism that leads to the tagging of PFC cell assemblies reported in contextual fear conditioning and that subsequently mature during the consolidation process (*52, 53*).

In the electrophysiological results we have shown that dendritic activity and plasticity in response to EC inputs is preserved during PV-cell activity manipulations (Fig. 1) and, therefore, the association of medial and lateral EC inputs in the dendrites of GCs in the behavioural experiments was likely not affected. This association, sometimes referred to as DG binding (*54*), is fundamentally different from the PV-operated DG binding mechanism that we introduced here. In our case, the binding refers to the association of multiple cell assemblies across brain regions. Both binding mechanisms, however, may contribute to pattern separation in the DG due to multiple feature binding, since the more exact and comprehensive the memory representation of an experience is, the easier it would be to discriminate from previous stored experiences with overlapping features. The contribution of systems-level binding to pattern separation was unveiled by the experiments in which we showed that GCs activation sparsity, a fundamental DG property for pattern separation (*2– 4*), was not affected by PV-cell inhibition, but context discrimination was however enhanced.

### Concluding remarks

We have presented imaging, electrophysiological and behavioural data that overall demonstrate a critical role of the DG in coordinating neuronal activity in a large network of brain structures known to be fundamental for memory formation. Using cell-specific manipulations we have shown that this systems-level gating mechanism can be operated by PV+ interneurons. We propose that through this simple mechanism, the DG fulfils two complementary functions, to select relevant information for updating memory, and to discard redundant information preserving the memory base from continuous overwriting. This mechanism merits further investigation in pathological conditions in which the proper updating of the memory base might be compromised. For instance, while a balanced increase in the inhibitory tone may prevent inefficient memory overwriting, an exacerbated or dysregulated DG inhibitory tone may render memories fixed, even after contingencies have changed, which might be potentially relevant for conditions such as Posttraumatic Stress Disorder (*55*). More generally, our results unveiled a new mechanism to control activity propagation in a complete network by regulating activity at a single node. Finding influential nodes in the brain network can help us develop methods for retuning maladaptive network dynamics (*13*).

## Materials and Methods

### Animals

All animal experiments were approved by the Animal Care and Use Committee of the Instituto de Neurociencias de Alicante (Alicante, Spain) and comply with the Spanish law (53/2013) and European regulations (EU directive 2010/63/EU).

In total, 143 mice, both males (n=82) and females (n=61), with two months of age at the beginning of the experiments, were randomly assigned to the different experimental groups (see below). Ten additional animals were used but excluded due to poor behavioural performance (less than 8 seconds of object exploration in the allocated 10 minutes of the novel object location task). No differences between sexes were found and data were pooled. Mice were bred in house from the line 129P2-Pvalb^tm1(cre)Arbr^/J (Jackson Laboratories, RRID: IMSR_JAX:008069) and housed in groups (3-5), with 12-12 h light/dark cycle, lights on at 8:00, at room temperature (23 ± 2 °C) and free access to food and water.

### Viral injections

#### Surgery

Mice were anaesthetized with isofluorane (Laboratorios Esteve, Murcia, Spain) 4%, 0.8L/min oxygen for induction, 1-2%, 0.8L/min oxygen for maintenance, and then fixed in a stereotaxic device (David Kopf Instruments, California, USA) over a heating pad at 37°C. Coordinates for targeting the injections in the hilus of the DG, from Bregma, were -2mm AP, ± 1.4 mm LM, +2mm DV (*56*). After opening the skin, we opened a 700µm Ø trepan with a milling cutter (Ref: FST 19007-07, Finest Science Tools, FST, Heidelberg, Germany) attached to a cordless micro drill (Ref: 58610V, Stoelting Co., Illinois, USA). Then we gently introduced a micropipette (Ref: 4878, World Precision Instruments, WPI, London, UK) through the trepan, into the brain. Micropipette was filled with mineral oil and viral vectors containing Designer Receptors Exclusively Activated by Designer Drugs, DREADDs (see below). Oil and viral vectors were separated with c.a. 1 µl of air. Micropipette was attached to a pump infusion Nanoliter 2010 Injector (WPI) coupled to the stereotaxic frame and was previously pulled in a P-2000 puller (Sutter Instruments Company, California, USA). We placed the tip of the micropipette in the hilus and very slowly injected 0.5 µl of viral vectors per hemisphere. After injection was completed, we waited 10 minutes before retracting the micropipette 200 µm, then waited another 10 minutes and finally removed gently and completely the pipette. After closing the skin with silk thread we administered subcutaneously analgesic, buprenorphine (Dechra Veterinaria Products SLU, Barcelona, Spain), and antibiotic, Syvaquinol (Laboratorios Syva S.A.U., León, Spain), to the mice and we controlled their recovery, injecting additional doses of analgesic if necessary.

### DREADDs, Designer Receptors Exclusively Activated by Designer Drugs

To modulate PV interneuron activity in the PV-cre mice we used Designer Receptors Exclusively Activated by Designer Drugs (DREADDs) (UNC Gene Therapy Vector Core, University of North Caroline, North Caroline, USA; Viral Vector Facility, Neuroscience Center Zurich, Switzerland). To inhibit PV cells we injected adeno-associated virus 5 (AAV5)-human synapsin1 (hSyn)-DIO-hM4D(Gi)-mCherry, and for activation we used the AAV5-hSyn-DIO-hM3D(Gi)-mCherry, as well as control virus synapsin1 (hSyn)-DIO-mCherry (*28*). The specificity of the expression in PV cells was confirmed immunohistologically (Supp. Fig. 1). After viral injection we allowed 3-4 weeks for DREADDs expression. Clozapine-N-Oxide (CNO, 1mg/kg, i.p in saline 0.9 %; ENZO Life Science Inc., New York, USA), was used for DREADDs activation (*28*). Behaviour

### Novel Object Location task (NOL)

After a period of handling, mice were introduced into an empty arena (50×50×30 cm) with spatial cues, and softly illuminated (luxes: 23 in the center ± 2 in the corners), and were allowed to freely explore it for 2 periods of 5 minutes (habituation phase). Twenty-four hours later we introduced two identical objects in the arena (located in opposite corners, 13’5 cm away from the walls) and the animals were allowed to explore them (familiarization or encoding phase). Familiarization with the objects was terminated when an animal reaches 20 seconds of accumulative exploration of both objects, or after 10 minutes in the arena (*30*). Animals exploring the objects less than 8 s in the 10 min of the familiarization phase were removed from the study (*a priori* exclusion criteria). Twenty-four hours later, animals entered in the same arena with the same objects, but one of them was displaced to a new location (another corner; 13’5 cm away from the walls and 15 cm away from the other object) and we again let the animals to explore under the same criteria (test or retrieval phase) (Supp. Fig. 2A and B). In one experiment (Fig. 2F), the object displacement was reduced to 10 cm to increase the task difficulty. We set the time that the animals explore the object in the new location divided by the total time of exploration of both objects, as an index of the mice spatial memory. Using this behavioural protocol, and injecting CNO or its vehicle in different moments, we could modulate the activity of PV interneurons during the encoding phase (injecting CNO or saline 90’ before the familiarization phase; n=38), during the consolidation (injecting CNO or saline 10 minutes after the familiarization phase; n=42) or during the retrieval (injecting CNO or saline 90’ before the test phase; n=41) (Fig. 2A). For encoding and control experiments, same animals were used. Different groups were used for consolidation and retrieval experiments.

In addition to the objects exploration, we measured and quantified locomotor parameters as movement velocity, distance travelled and side vs centre arena preference (Supp. Fig. 2B and C).

### Elevated plus maze (EPM)

We tested potential changes in anxiety induced by the modulation of PV-cell activity in the DG, using the elevated plus maze (EPM). One week after the NOL task, animals were injected either with CNO or saline, and placed 90’ later in a EPM with two open and two closed arms (Supp. Fig. 2D). Animals were video-recorded and its performance tracked with commercial software (Smart Video Tracking Software, Panlab, Barcelona, Spain). We compared the time that the animal spent in open *vs*. closed arms as an index of anxiety. We also checked for other parameters as number of entrances in each arm and distance travelled (Supp. Fig. 2E).

After the behavioural evaluation, mice were divided in groups for further exploration in electrophysiological or fMRI experiments.

### Electrophysiology

Mice were anaesthetized with urethane (1.4 g/kg, i.p.) and recording and stimulating electrodes were implanted following stereotaxic standard procedures. We introduced one bipolar stimulating electrode (WPI, ref. TM53CCNON) in the perforant pathway (from bregma: -4.3 AP, +2.5 ML, +1.4 DV, -12°; DV position was adjusted per animal to optimize the evoked response), and two recording probes (single shank, 50 µm contact spacing, 32 channels; NeuroNexus Technologies, Michigan, USA) were targeted to the PFC (+2 AP, +0.2 ML, +2.5 DV) and hippocampus (−2 AP, +1.5 ML, -2 DV), including CA1 and DG regions. Recordings were performed before and up to 3 h after CNO injection (1mg/kg, i.p.). We induce LTP using a standard theta burst protocol (six 400 Hz-trains of pulses delivered at a 200 ms interval, repeated six times at an interval of 20 s), and 60 min after induction we recorded again evoked potentials. After filtering (0.1-3k Hz) and amplifying, the electrophysiological signals were digitalized (20-32 kHz acquisition rate) and the data analysed offline using the software Spike2 (Cambridge Electronic, Cambridge, UK) or MatLab (The MathWorks Inc., Natick, MA, USA). We measured the excitatory postsynaptic potential (EPSP) as the maximum slope of the evoked potential in the molecular layer of the DG and in the *stratum radiatum* of CA1, and the population spike (PS) as the amplitude of the fast negative potential recorded in hilus of the DG and in the CA1 pyramidal cell layer. In PV-Gi animals, in which CNO administration induced a large increase in the PS amplitude, the EPSP was measured more distal from the granule cell soma layer (in the outer third of the molecular layer), giving smaller estimations. This is a common procedure to avoid the volume conducted artefact of the PS that would have prevented the precise measure of the EPSP slope.

### Functional MRI

For the fMRI experiments, mice were anaesthetized with 1.4 g/kg of urethane and implanted with custom made carbon fibre MRI-compatible electrodes as described previously (*44*). Briefly, individual 7 µm diameter carbon fibres (Goodfellow Cambridge Limited, Cambridge, UK) were bundled and introduced into a pulled double borosilicate capillary (WPI, ref. TST150-6”) with ≃ 200 µm tip diameter and electrical impedance of 40-65 kΩ. Tip was bent in a flame to form a 90° angle to minimize the distance between the head of the animal and the magnetic resonance imaging array coil once implanted. A regular wire connector was couple to the pipette, connected to the carbon fibres using silver conductive epoxy resin (Ref. 186-3616, RS components, Madrid, Spain), and isolated with rapid epoxy resin (Araldite, Basel, Switzerland). The electrode was then implanted in the perforant pathway of the mice (same coordinates than previously), and its optimization guided by the evoked potential recorded in DG (same methods than for *in vivo* electrophysiology). Once optimized, we removed gently the recording electrode to avoid any bleeding and the carbon fibre electrode was secured to the skull with super-bond C&B dental cement (Sun Medical Co. LTD, Moriyana, Shiga, Japan). Then mice were placed in a custom made animal holder with adjustable bite and ear bars and positioned into a horizontal 7-T scanner (Biospec 70/30, Bruker Medical, Ettlingen, Germany). Animals were constantly supplied with 0.8 L/min O_2_ and temperature was controlled between 36.5 and 37.5 °C with a water heat-pad.

Functional imaging acquisition was performed in 12 coronal slices using a GE-EPI applying the following parameters: field of view (FOV) 25×25 mm, slice thickness, 1 mm; matrix, 96×96; segments, 1; FA 60°; time echo (TE), 15 ms; time repetition (TR), 2000 ms. T_2_-weigthed anatomical images were collected using a rapid acquisition relaxation enhanced sequence (RARE): FOV, 25×25 mm; 12 slices; slice thickness, 1mm; matrix, 192×192; TE_eff_, 56 ms; TR, 2 s; RARE factor, 8. A 1H mouse brain received –only phased-array coil with integrated combiner and preamplifier, and no tune/no match, was employed in combination with the actively detuned transmit-only resonator (Bruker BioSpin MRI GmbH, Germany).

Functional MRI data were analysed offline using our own software developed in MatLab (The MathWorks Inc.), which included Statistical Parametric Mapping packages (SPM8, www.fil.ion.ucl.ac.uk/spm). After linear detrending, temporal (0.015-0.2 Hz) and spatial filtering (3×3 Gaussians kernel or 1.5 sigma) of voxel time series, a general linear model (GLM) was applied. Functional maps were generated from voxels that had a high significant component for the model (*p* < 0.01) and were clustered together in space. ROIs were extracted from mice brain atlas (*56*) and included hippocampus, PFC (prelimbic, infralimbic, orbitofrontal and cingular), medial temporal lobe structures (entorhinal cortex, subiculum, presubiculum, perirhinal and ectorhinal cortex), subcortical structures (amygdala, basal nuclei and NAc) and cortical areas (visual cortex, auditory cortex, sensory cortex, associative areas and motor cortex). We calculate the number of active voxels before and after CNO injection (1 mg/kg, i.p.) when stimulating at 10Hz or 20 Hz in the performant pathway (4 s ON, 26 s OFF).

After electrophysiology or fMRI experiments, animals were perfused and their brain extracted for histology.

### Histology

Animals were perfused with 50 ml of 37°C saline and 50 ml of 4% paraformaldehyde (PFA; BDH prolabo, VWR chemicals, Lovaina, Belgium). Brains were extracted and kept in PFA 4% for at least 24h at 4°C and then cut in a fixed material vibratome (VT 1000S, Leica, Wetzlar, Germany) in 50 µm slices. Validation of anatomical coordinates for recording and stimulating electrodes was done in DAPI stained slices. Viral infection efficiency and specificity were quantified in 6-8 hippocampal containing slices following standard immunohistochemistry protocols for PV (primary antibody: mouse anti-PV, 1:2000, ref. 235, Swant, Switzerland; secondary antibody: Alexa 488 goat anti-mouse, 1:500, ref. A11029, Life Technologies; USA) and using the virus reporter mCherry co-labelling.

### c-Fos experiments

After a period of handling, a total of 22 animals (8 PV-Sham, 8 PV-Gi, 6 PV-Gq) were placed in the NOL arena for the encoding phase, and sacrificed 60 min later. Perfusion of the animals and tissue slicing was performed as indicated above. Eight to ten serial slices covering the hippocampus, PFC and NAc were c-Fos immunostained (primary antibody: guinea pig anti-cFos, 1:1000, ref. 226004, Synaptic Systems, Göttinger, Germany; secondary antibody: Alexa 488 donkey anti-guinea pig, 1:500, ref. 706-545-148, Jackson ImmunoResearch, Suffolk, United Kingdom) mounted and used in the quantitative analysis (below).

Quantitative analysis of c-Fos positive nuclei was performed offline on 12-bit grey scale images acquired in a Leica DM4000 fluorescence microscope at 10x/0.25 dry objective using a Neurolucida software (MBF Bioscience, Williston, VT USA). The ROIs were manually delineated following the Franklin and Paxinos mouse brain atlas (*56*). Analysis were performed using Icy Software (*57*) in a semi-automatic manner. The threshold for detection of positive nuclei was set for each brain region, setting average nuclei size and a signal/noise ratio higher than 23%, according to Rayleigh criterion for resolution and discrimination between two points. Animals with misplaced viral infections were removed from all the analysis.

### Statistical analysis

The statistical analysis was done using GraphPad Prism 7 software (GraphPad Software Inc., La Jolla, CA, USA) or SPSS v20 software (IBM, New York, USA). After an exploratory analysis for the presence of outlier values, we checked the skewness and kurtosis of the data before testing their statistical distribution. Parametric-test requirements, including normality (D’Agostino-Pearson test and Shapiro-Wilk test) and homoscedasticity (F of Levene test) were tested. All the data fulfilled parametric criteria, unless otherwise specified. In most analysis we applied two-way ANOVA, with a *group* factor with 3 levels (Sham, PV-Gi and PV-Gq), and a *time* factor with 2 levels (before and after CNO injection). In case of c-Fos analysis, we applied one-way ANOVA with group as factor with the same 3 levels as above. We applied a Sidak *post hoc* analysis for multiple comparisons with adjusted alpha. Effect size was calculated with partial *eta* square (η_p_^2^) as SSeffect / (SSeffect +SSerror), and the power of the effect (1-β) using GPower software (University of Düsseldorf, Düsseldorf,

Germany). Effect size is considered large if η _p_ ^2^ > 0.13. If, by any reason, partial *eta* square could not be used, then we express r^2^ indicator, being similar to *eta*, but requiring > 0.30 values to indicate large effects. The power of the effect (1-β) is taken as an invert indicator of committing a type II error. The results of the statistical analysis are detailed in Supp. Tables numbered according to the number of the Figures to which they correspond.

For regression analysis, Pearson’s correlation was applied and coefficients were transformed to apply Fisher’s 1925 test (*58*) for significant values, using RStudio and *cocor* package (RStudio 2015. Inc., Boston, MA). To test the importance of DG firing coordination over the signal propagation on CA1 before vs after CNO modulation we did a General Linear Model analysis of CA1 PS including the width of the DG PS in its half amplitude as a between groups co-variable.

Data were plotted using GraphPad Prism 7 software (GraphPad Software Inc.) and figures were composed using Adobe Illustrator software (Adobe Systems Incorporated, San José, California, USA).

## Acknowledgments

We thank Begoña Fernández, Jesús Pacheco and Víctor Rodríguez for excellent technical assistance. This work was supported by the Spanish State Research Agency through the Severo Ochoa Program for Centres of Excellence in R&D (SEV-2017-0723) and the Ministerio de Economía y Competitividad (MINECO) and FEDER funds under Grant No. BFU2015-64380-C2-1-R and PGC2018-101055-B-I00. J.M.C was supported by a FPU fellowship from the Ministerio de Educación of Spain (ref. 2012/1151); E.P-M was supported by a fellowship from the Consellería de Educación, Invetigación, Cultura y Deporte (ref. ACIF/2016/111).

## Authors contributions

SC conceptualized the project; J.M.C., E.P.-M. and R.G.-H. performed investigation; All authors analysed the data; J.M.C. and SC wrote the original manuscript and all the authors reviewed and edited the final manuscript, SC acquired funds and supervised research.

## Competing interests

Authors declare no competing interests.

## Supplementary Materials

### Supplementary figures

**Supp. Fig. 1.**
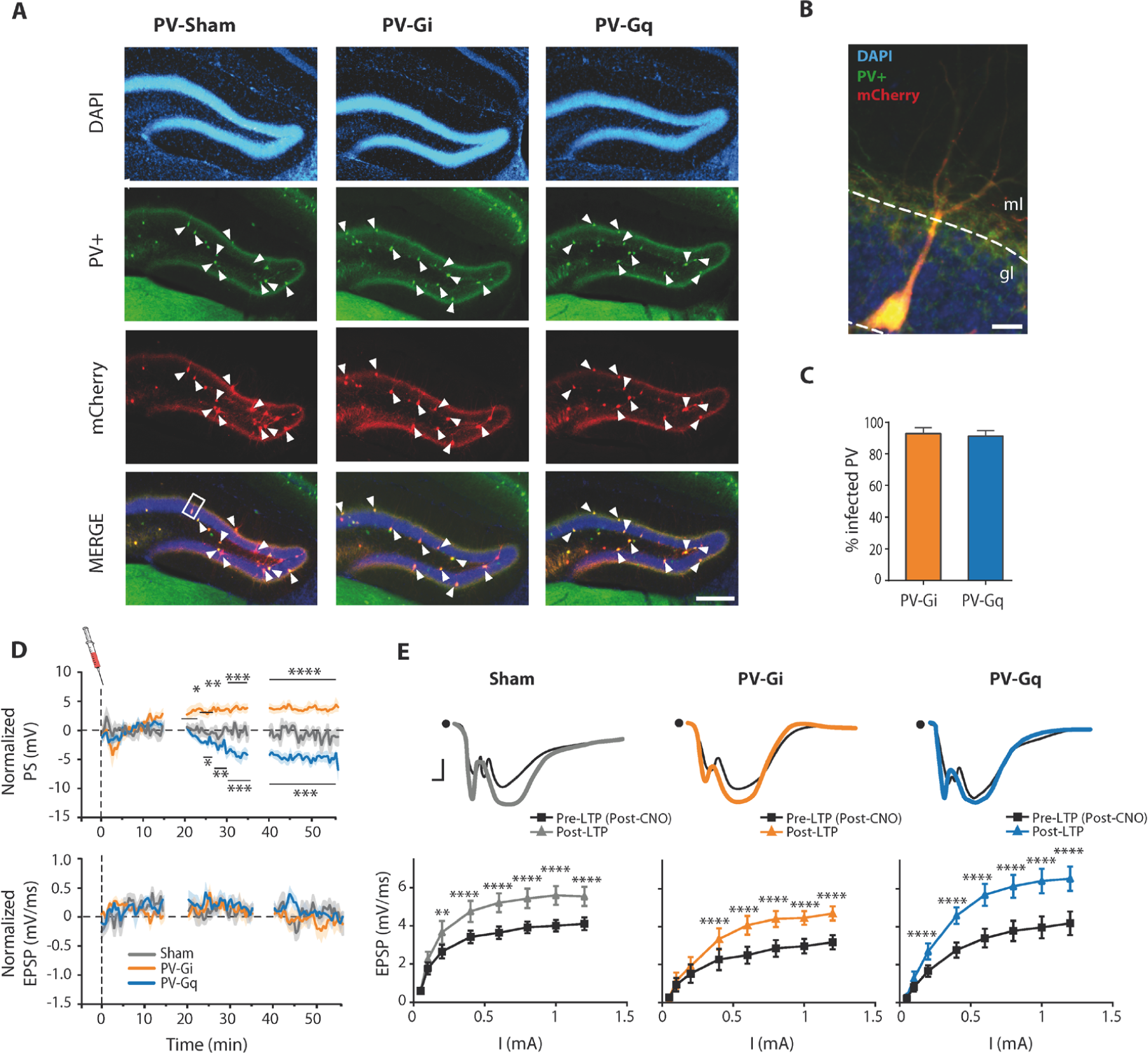
Expression of DREADDs in the DG of PV-cre animals and modulation of granule cell activity. (Complements main Fig. 1) **A**, Representative pictures showing the reporter of the virus infection (mCherry, red), immunolabelled against PV protein (green) and counterstained with DAPI (cell nuclei, blue). White arrows point to some co-localization examples. Scale bar: 100 µm. **B**, Zoom in of an infected PV+ cell (white square in A). Scale bar: 10 µm. **C**, Efficiency of DREADDs expression in the DG expressed as the % of infected PV-cells in PV-Gi (yellow) and PV-Gq (blue) animals. Data show mean ± SEM. **D**, Temporal course of the evoked DG PS amplitude (top) and EPSP slope (bottom) after CNO injection (1 mg/kg, i.p.) in response to single pulse stimulation of the PP. The first hour of the c.a. 3h of each recording session is shown. Solid line and shadow represent the mean and SEM, respectively. **E**, Synaptic potentiation in all animal groups. LTP was induced 1 h after CNO administration in all groups. Shown are the EPSP waveforms (upper panels) and stimulus response curves (lower) recorded immediately before (pre-) and 1 h after (post-) LTP induction. Representative evoked responses are shown in the upper part. Note that the overall smaller EPSP slope in the PV-Gi group is the consequence of measuring the waveform in positions more distal to the granule cell soma (see Fig. 1 and Methods Section). This was done to avoid the volume conducted artefact introduced by the substantially larger and faster PS in this group. All statistical values are detailed in Supp. Table 5. *p≤0.05, **p≤0.01, ***p≤0.001, ****p≤0.0001.

**Supp. Fig. 2.**
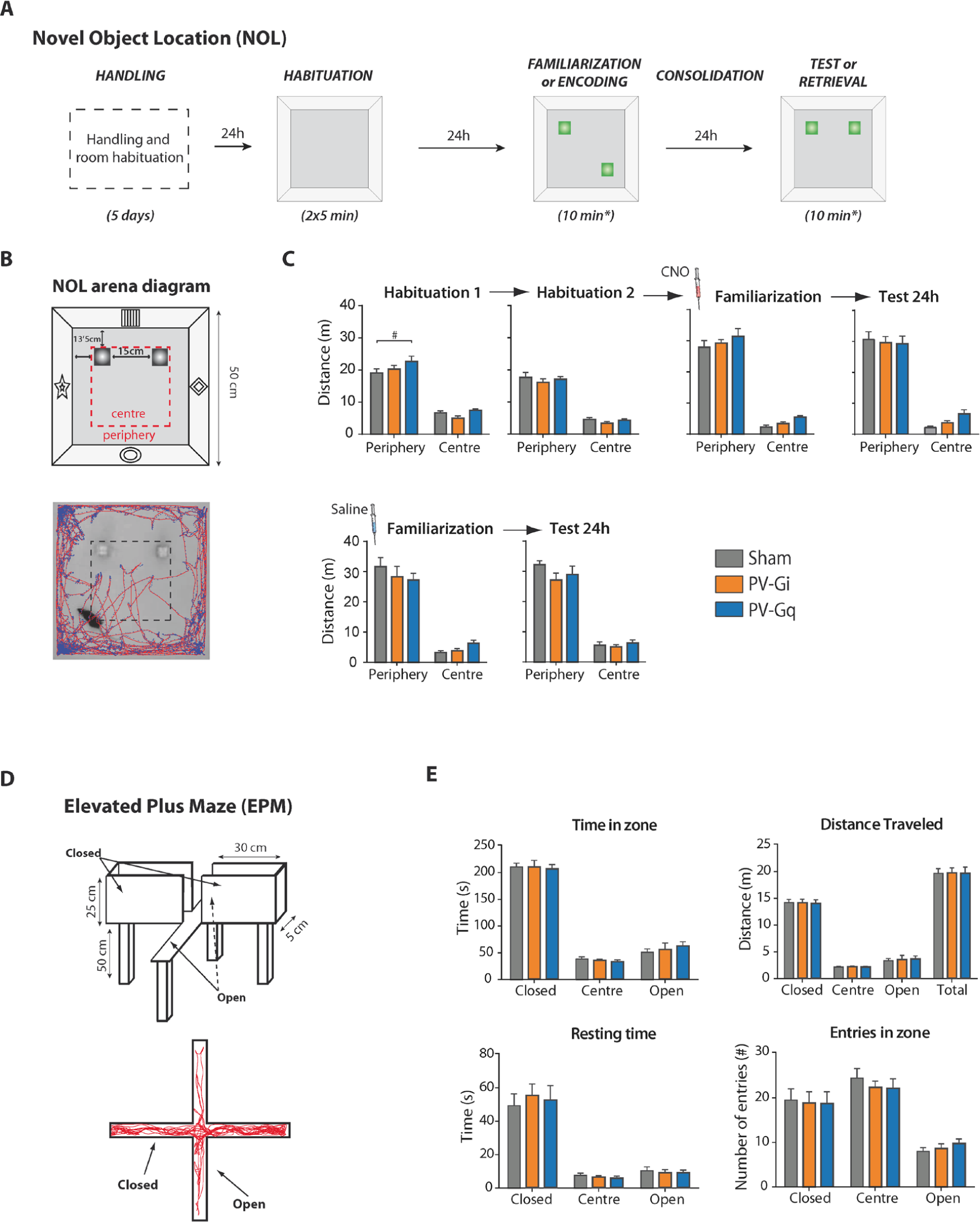
Motor activity and anxiety. (Complements main Fig. 2) **A**, Diagram of the Novel Object Location (NOL) protocol followed. **B**, Detailed diagram of the NOL arena. Example of one mouse tracking within a test session (bottom). **C**, For evaluating locomotor activity, with and without PV modulation, the travelled distance in each session was analyzed for the three groups of mice. **D**, Diagram of the Elevated Plus Maze used for evaluating anxiety (top). Example of one trial (bottom). **E**, Anxiety levels as measured by time spent, distance travelled, resting time and entries per zone. All statistical values are detailed in Supp. Table 5.

**Supp. Fig. 3.**
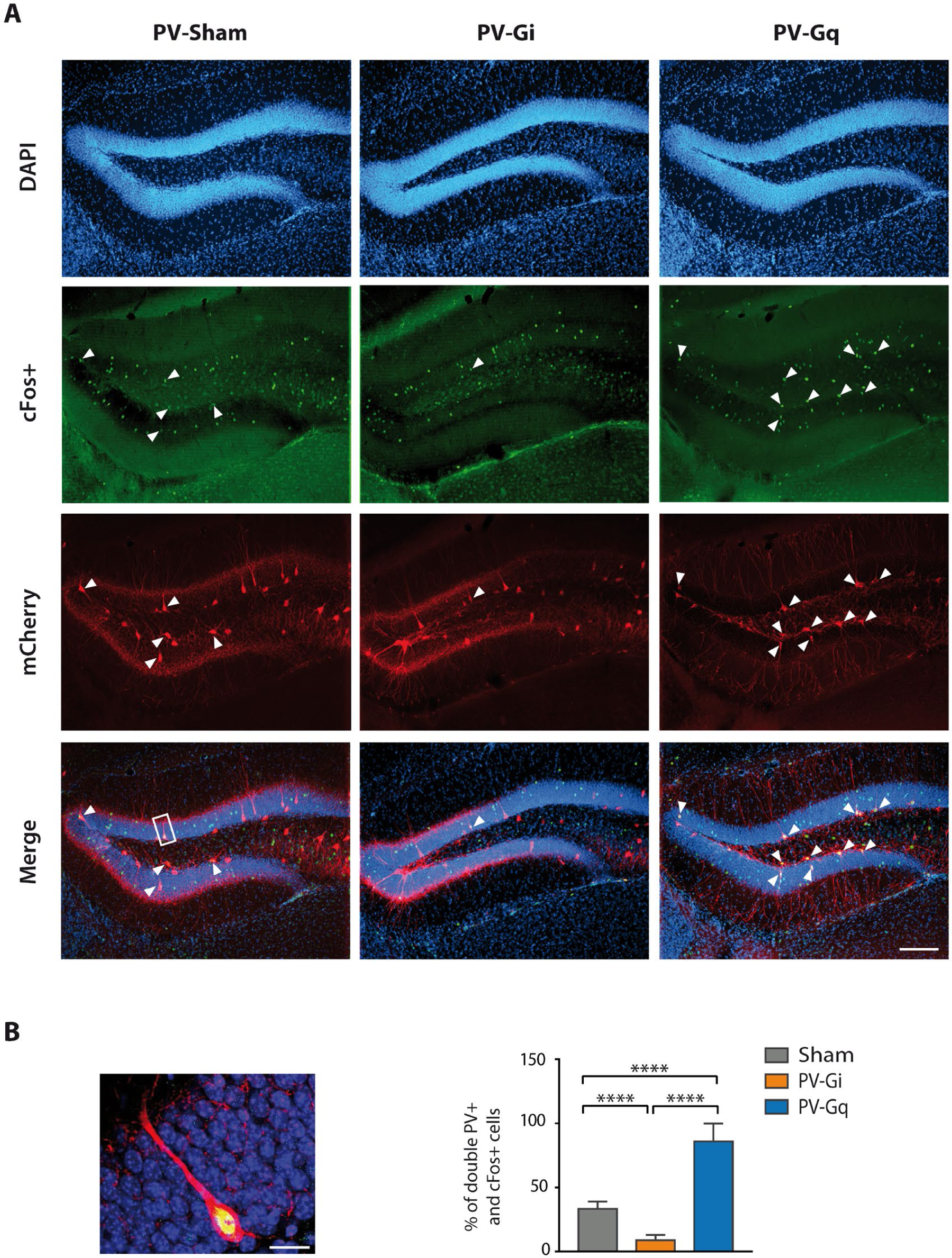
c-Fos expression in the hippocampus induced by context exploration. (Complements main Fig. 2) **A**, Representative microphotographs of the DG showing c-Fos+ cells (green), infected PV-cells (mCherry, red) and their co-localization (merge). Scale bar: 100 µm. **C**, Zoom in of a double PV+ and c-Fos+ cell (from the white rectangle in **A**), and their quantification in all groups. Data represent mean ± SEM. All statistical values are detailed in Supp. Table 5. ****p≤0.0001.

**Supp. Fig. 4.**
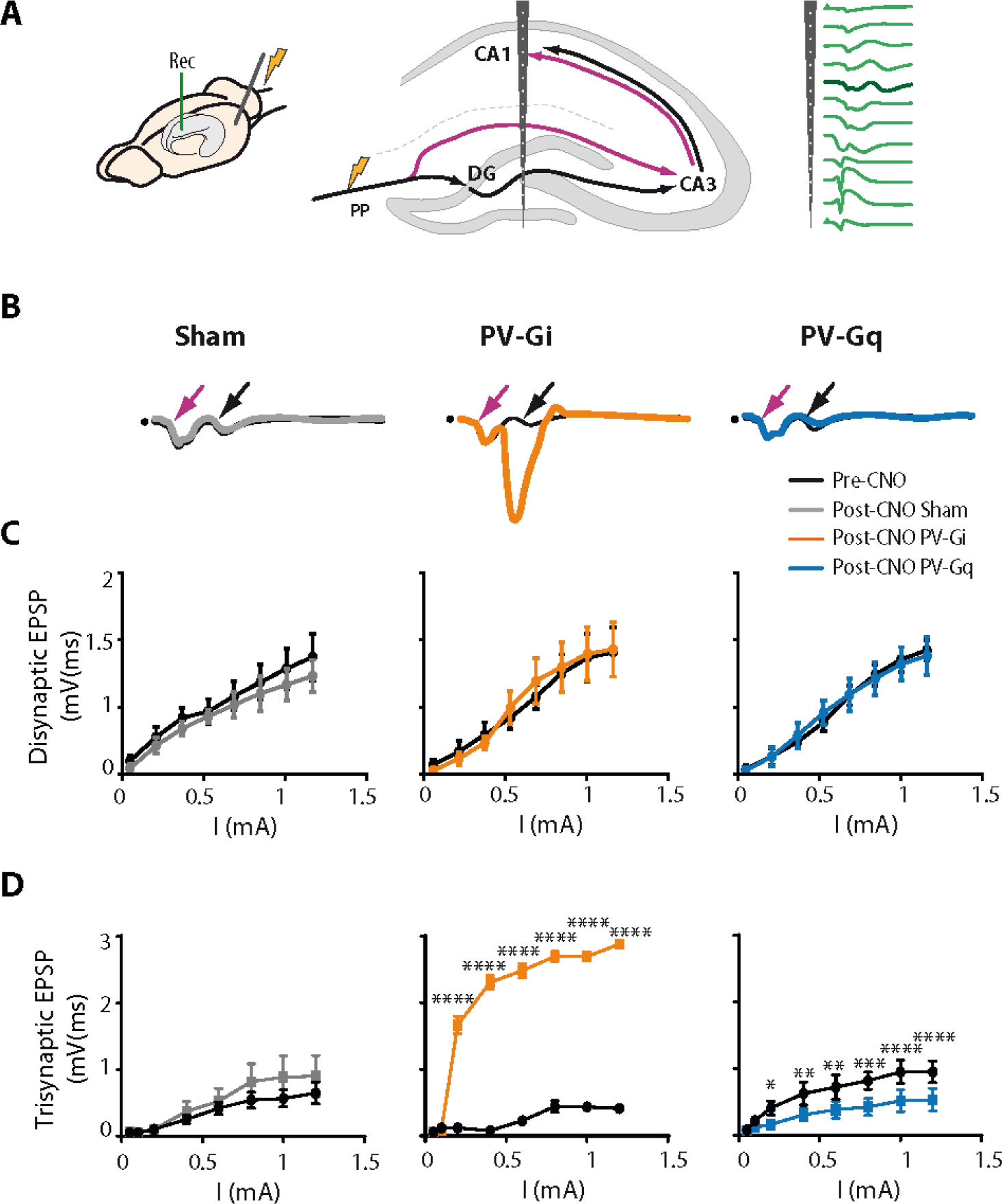
Dentate gyrus PV+ interneurons regulate activity propagation in the hippocampus strongly and specifically in the tri-synaptic circuit. (Complements main Fig. 3) **A**, Schematic representation of the trisynaptic (PP→DG→CA3→CA1; black) and disynaptic (PP→CA3→CA1; purple) hippocampal circuits and representative multichannel *in vivo* electrophysiological recordings (green traces) in response to PP stimulation. Thick trace indicates the selected channel for the analysis of evoked di- and tri-synaptic EPSPs. **B**, Representative waveforms recorded in CA1 *stratum radiatum* in response to a PP stimulation. Purple and black arrows points to di-synaptic and tri-synaptic EPSPs, respectively. Scale bars: 4ms and 0.5 mV. **C**, Comparison of PP stimulus-response curves of CA1 di-synaptic EPSP slope before (black) and after CNO i.p. injection in Sham (grey), PV-Gi (yellow) and PV-Gq (blue) animals. **D**, Same as (**C**) but for the tri-synaptic CA1 EPSP. Group data represents mean ± SEM. All statistical values are detailed in Supp. Table 5. *p≤0.05, **p≤0.01, ***p≤0.001, ****p≤0.0001.

**Supp. Fig. 5.**
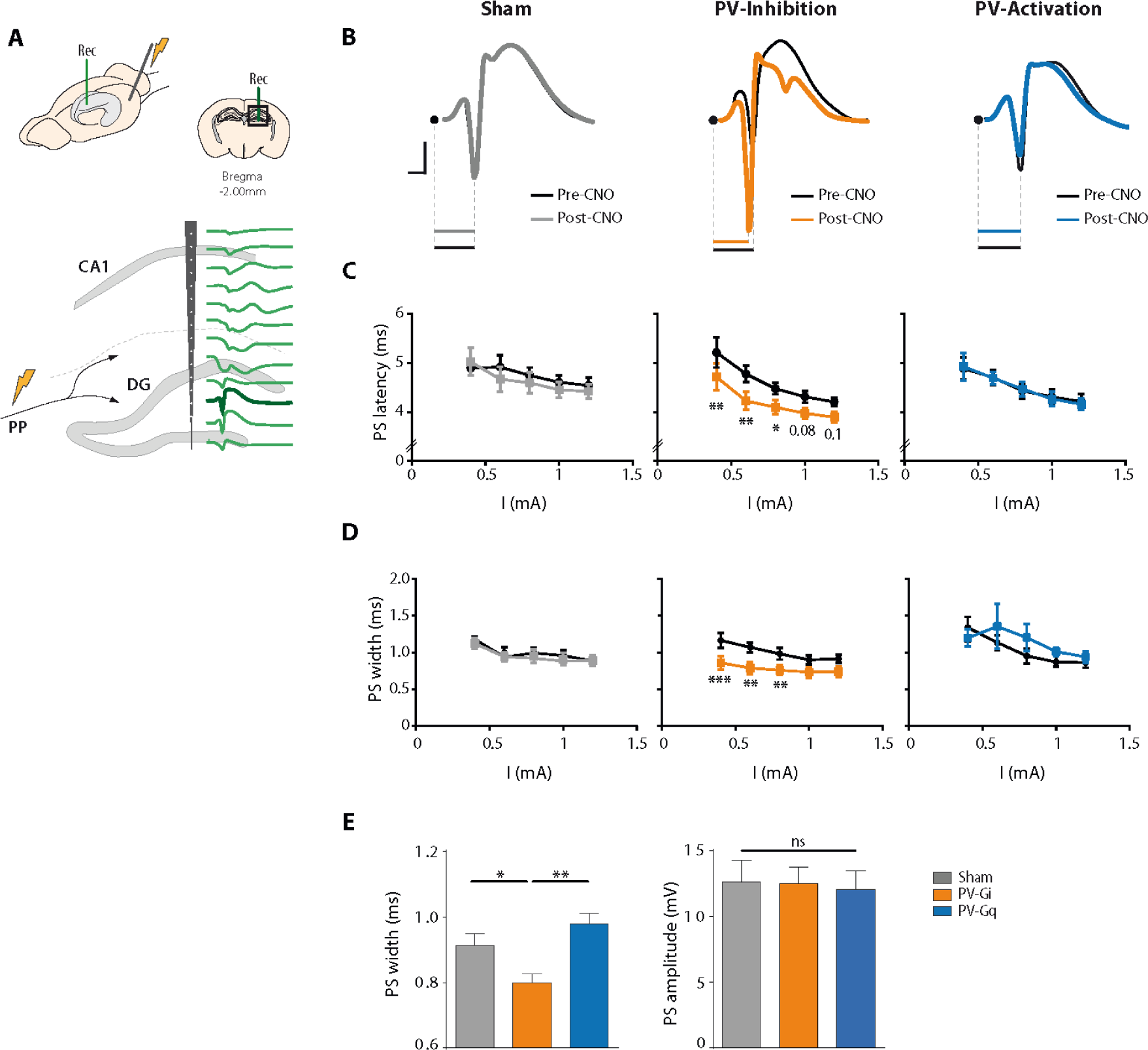
Firing delay and synchrony of granule cells. **A**, Schematic representation of multichannel in vivo electrophysiological recordings (green traces) in response to PP stimulation. Thick trace indicates the selected channel for the analysis of the PS. **B**, Representative waveforms of DG PS, highlighting the differences in firing latency. Stimulation artefact is replaced by a black dot. **C**, Stimulus-response curves for the PS latency in the DG. **D**, Firing synchrony of granule cells as measured by the width (at half maximum amplitude) of the PS (higher synchrony implies smaller width). **E**, Higher synchrony (smaller width) is not the consequence of larger PS amplitudes. Comparably sized PS result in significantly different widths across experimental groups. Group data represents mean ± SEM. All statistical values are detailed in Supp. Table 5. *p≤0.05, **p≤0.01, ***p≤0.001.

**Supp. Fig. 6.**
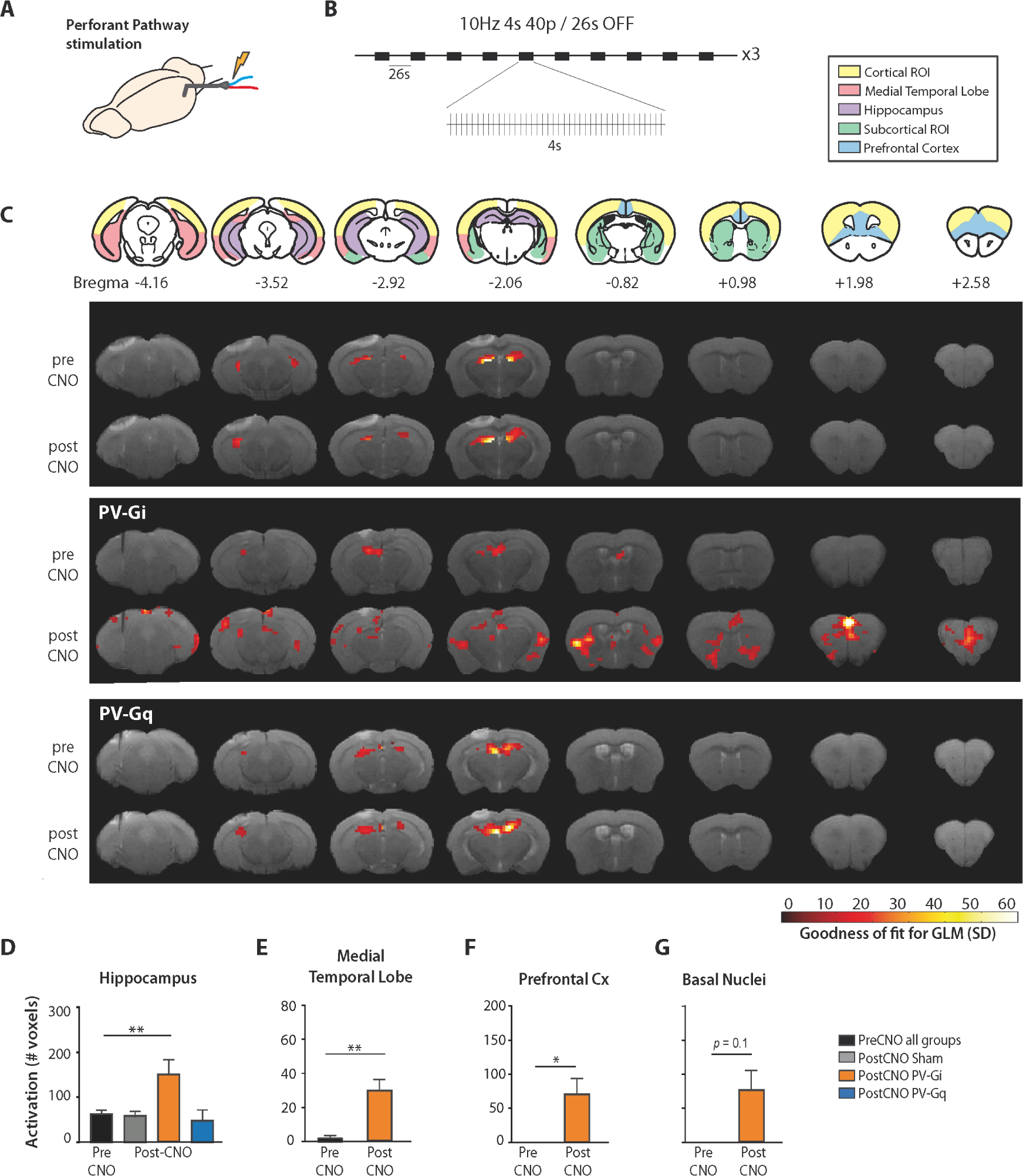
Brain-wide functional connectivity regulated by DG PV-cell activity. (Complements main Fig. 4) **A**, Schematic of the preparation. **B**, 10 Hz stimulation protocol (40 pulses per train) applied in the PP. **c**, Upper: regions of Interest (ROIs) used in the quantitative analysis. Numbers indicate mm from Bregma. Lower: functional maps evoked by the stimulation before and after CNO injection, and overlaid on T2 anatomical images. Colour-code represents the goodness of fit of the GLM analyses thresholded at p<0.01. **D-G**, Number (mean ± SEM) of statistically significant active voxels per group and condition in the indicated ROIs. Pre-CNO data are represented together for simplicity. All statistical values are detailed in Supp. Table 5. *p≤0.05, **p≤0.01.

**Supp. Fig. 7.**
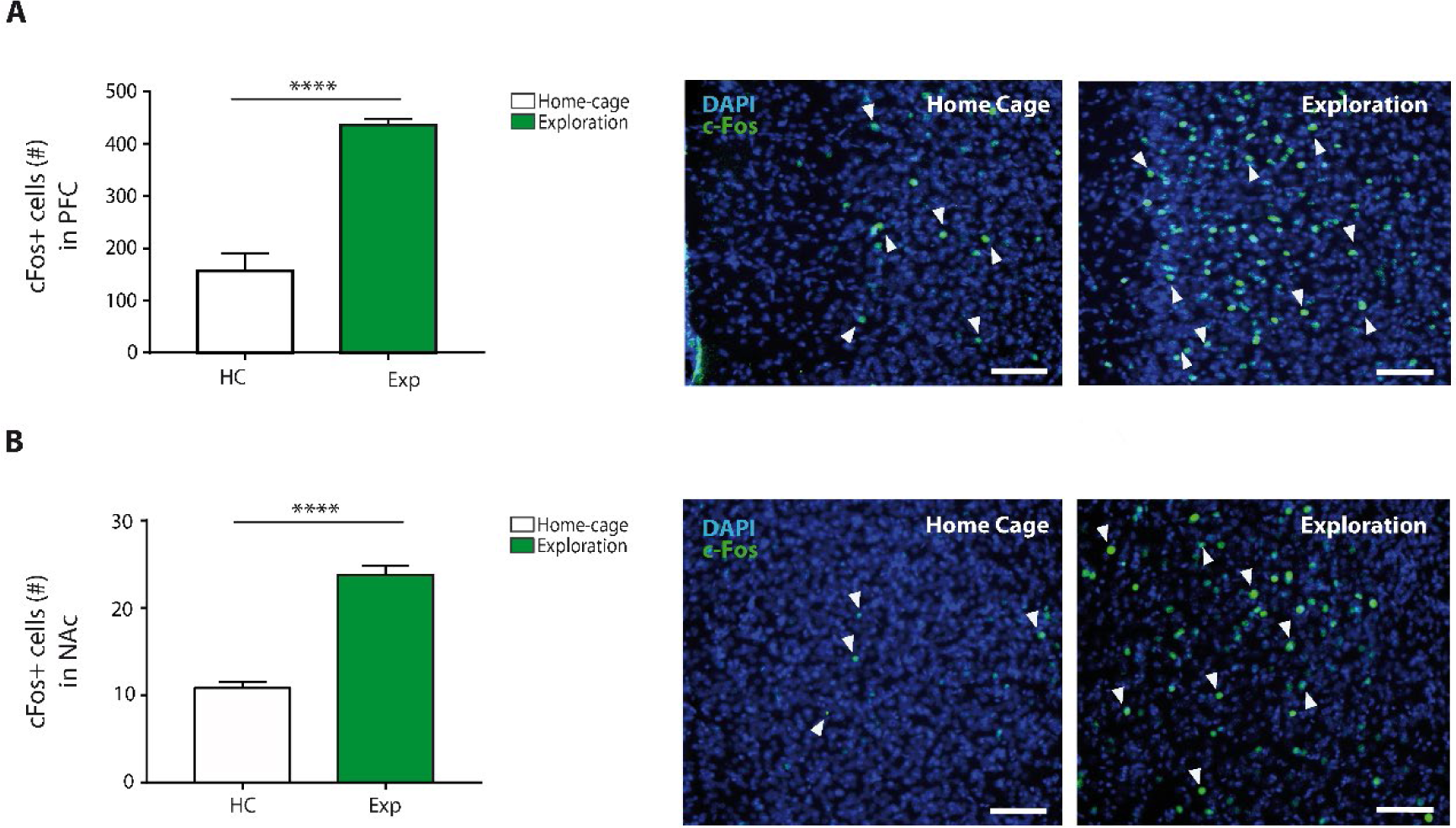
c-Fos expression in PFC and NAc during NOL encoding phase *vs*. homecage. (Complements main Fig. 2) **A**, Number of c-Fos+ cells in the medial prefrontal cortex (PFC) activated by exploration during the familiarization (encoding) phase of the NOL task (green) compared to the baseline activity in the hime cage (white). Insets: representative c-Fos immunostained cells (white arrows) in the PFC for both groups. **B**, Same for nucleus accumbens (NAc). Scale Bars: 100 μm. All statistical values are deailed in Supp. Table 5. ****p≤0.0001.

### Supplementary Tables

**Supp. Table 1.** Statistical data corresponding to Figure 1, including the statistical test, sample size (n), value of the statistic, degrees of freedom (df), associated *p* value, effect size (η _p_ ^2^) and power effect (1-β).

| Panel | Analysis/group | Statistic | Value | df | *p* value | $\eta_p^2$ | 1-β |
| --- | --- | --- | --- | --- | --- | --- | --- |
| e | Sham (n=6) | Two ways ANOVA, RM | F = 1.16 | 7,40 | 0.16 | - | - |
|  | PV-Gi (n=6) |  | F = 3.97 | 7,40 | 0.002* | 0.41 | 0.9 |
|  | PV-Gq (n=6) |  | F = 4.81 | 7,40 | 0.0005* | 0.45 | 0.94 |
| f | Sham (n=6) | Two ways ANOVA, RM | F = 0.56 | 7,40 | 0.78 | - | - |
|  | PV-Gi (n=6) |  | F = 0.21 | 7,40 | 0.98 | - | - |
|  | PV-Gq (n=6) |  | F = 0.03 | 7,40 | 0.99 | - | - |
| i | Between groups, PreLTP | Two ways ANOVA | F = 2.02 | 2,60 | 0.14 | - | - |
|  | Between groups, PostLTP | Two ways ANOVA | F = 1.56 | 2,60 | 0.22 | - | - |
|  | Within group, Pre-PostLTP | Two ways ANOVA, RM | F = 162.5 | 5,30 | <0.0001* | 0.84 | 1 |
|  | Sham (n=6) | *post hoc*<sup>⓪</sup> | t = 15.06 | 30 | <0.0001* |  |  |
|  | PV-Gi (n=6) | *post hoc*<sup>⓪</sup> | t = 16.65 | 30 | <0.0001* |  |  |
|  | PV-Gq (n=6) | *post hoc*<sup>⓪</sup> | t = 17.38 | 30 | <0.0001* |  |  |

\* Statistically significant value.
<sup>⓪</sup> Sidak post hoc comparison, *alfa* adjusted by Bonferroni at 0.05 significance level.

**Supp. Table 2.** Statistical data corresponding to Figure 2, including the statistical test, sample size (n), value of the statistic, degrees of freedom (df), associated *p* value, effect size (η _p_ ^2^) and power effect (1-β).

| Panel | Analysis/group | Statistic | Value | df | p value | $\eta_p^2$ | 1-β |
| --- | --- | --- | --- | --- | --- | --- | --- |
| b | Between groups, PreCNO | One way ANOVA | F = 0.06 | 2,35 | 0.94 | - | - |
|  | Between groups, PostCNO | Two ways ANOVA, RM | F = 12.31 | 2,35 | <0.0001* | 0.41 | 1 |
|  | Sham vs PV-Gi | *post hoc*<sup>o</sup> | t = 3.29 | 70 | 0.004* |  |  |
|  | PV-Gi vs PV-Gq | *post hoc*<sup>o</sup> | t = 5.88 | 70 | <0.0001* |  |  |
|  | Sham vs PV-Gq | *post hoc*<sup>o</sup> | t = 2.65 | 70 | 0.02* |  |  |
|  | Within groups, Pre-PostCNO | Two ways ANOVA, RM | F = 55.86 | 1,35 | <0.0001* | 0.61 | 1 |
|  | Sham (n=13) | *post hoc*<sup>o</sup> | t = 4.27 | 35 | 0.0004* |  |  |
|  | PV-Gi (n=13) | *post hoc*<sup>o</sup> | t = 8 | 35 | <0.0001* |  |  |
|  | PV-Gq (n=12) | *post hoc*<sup>o</sup> | t = 0.81 | 35 | 0.8 |  |  |
| c | Between groups, PreCNO | One way ANOVA | F = 2.24 | 2,39 | 0.12 | - | - |
|  | Between groups, PostCNO | Two ways ANOVA, RM | F = 0.04 | 2,39 | 0.95 | - | - |
|  | Within groups, Pre-PostCNO | Two ways ANOVA, RM | F = 60.23 | 1,39 | <0.0001* | 0.6 | 1 |
|  | Sham (n=10) | *post hoc*<sup>o</sup> | t = 4.49 | 39 | 0.0002* |  |  |
|  | PV-Gi (n=17) | *post hoc*<sup>o</sup> | t = 5.14 | 39 | <0.0001* |  |  |
|  | PV-Gq (n=12) | *post hoc*<sup>o</sup> | t = 3.95 | 39 | 0.001* |  |  |
| d | Between groups, PreCNO | One way ANOVA | F = 0.89 | 2,38 | 0.41 | - | - |
|  | Between groups, PostCNO | Two ways ANOVA, RM | F = 0.74 | 2,38 | 0.48 | - | - |
|  | Within groups, Pre-PostCNO | Two ways ANOVA, RM | F = 38.16 | 1,38 | <0.0001* | 0.5 | 1 |
|  | Sham (n=14) | *post hoc*<sup>o</sup> | t = 4.27 | 38 | 0.0004* |  |  |
|  | PV-Gi (n=15) | *post hoc*<sup>o</sup> | t = 3.95 | 38 | 0.001* |  |  |
|  | PV-Gq (n=16) | *post hoc*<sup>o</sup> | t = 2.51 | 38 | 0.04* |  |  |
| e | Between groups, PreCNO | One way ANOVA | F = 2.35 | 2,35 | 0.11 | - | - |
|  | Between groups, PostCNO | Two ways ANOVA, RM | F = 0.28 | 2,35 | 0.75 | - | - |
|  | Within groups, Pre-PostCNO | Two ways ANOVA, RM | F = 29.28 | 1,35 | <0.0001* | 0.31 | 1 |
|  | Sham (n=13) | *post hoc*<sup>o</sup> | t = 2.88 | 35 | 0.019* |  |  |
|  | PV-Gi (n=13) | *post hoc*<sup>o</sup> | t = 2.81 | 35 | 0.023* |  |  |
|  | PV-Gq (n=12) | *post hoc*<sup>o</sup> | t = 3.65 | 35 | 0.0026* |  |  |
| f | Between groups (n=8) | Two ways ANOVA, RM | F = 6.11 | 2,28 | 0.0063* | 0.30 | 0.96 |
|  | Fam | *post hoc*<sup>o</sup> | t = 0.83 | 42 | 0.8 |  |  |
|  | 10 cm | *post hoc*<sup>o</sup> | t = 3.75 | 42 | 0.0016* |  |  |
|  | 15 cm | *post hoc*<sup>o</sup> | t = 0.02 | 42 | 0.99 |  |  |
|  | Within groups (n=8) | Two ways ANOVA, RM | F = 16.35 | 2,28 | <0.0001* | 0.56 | 0.99 |
|  | Saline fam vs 10 cm | *post hoc*<sup>o</sup> | t = 0.2 | 28 | 0.99 |  |  |
|  | Saline fam vs 15 cm | *post hoc*<sup>o</sup> | t = 3.57 | 28 | 0.004* |  |  |
|  | Saline 10 cm vs 15 cm | *post hoc*<sup>o</sup> | t = 3.36 | 28 | 0.007* |  |  |
|  | CNO fam vs 10 cm | *post hoc*<sup>o</sup> | t = 4.85 | 28 | <0.0001* |  |  |
|  | CNO fam vs 15 cm | *post hoc*<sup>o</sup> | t = 4.43 | 28 | 0.0004* |  |  |
|  | CNO 10 cm vs 15 cm | *post hoc*<sup>o</sup> | t = 0.42 | 28 | 0.97 |  |  |
| g | Home-cage vs Exploration<br>(n=2) (n=8) | t-test independent measures | t = 3.41 | 8 | 0.009 * | 0.59 | 0.1 |
| h | Sham, PV-Gi, PV-Gq<br>(n=8) (n=9) (n=6) | One way ANOVA | F = 1.93 | 2,20 | 0.17 | - | - |
| i | Sham, PV-Gi, PV-Gq<br>(n=8) (n=9) (n=6) | One way ANOVA | F = 0.3 | 2,20 | 0.74 | - | - |
| j | Sham, PV-Gi, PV-Gq<br>(n=8) (n=9) (n=6) | One way ANOVA | F = 0.91 | 2,22 | 0.41 | - | - |
| k | Between groups | One way ANOVA | F = 6.92 | 2,18 | 0.0059* | 0.3 | 0.7 |
|  | Sham (n=7) vs PV-Gi | *post hoc*<sup>o</sup> | t = 1.95 | 18 | 0.18 |  |  |
|  | Sham vs PV-Gq (n=5) | *post hoc*<sup>o</sup> | t = 1.96 | 18 | 0.18 |  |  |
|  | PV-Gi (n=6) vs PV-Gq | *post hoc*<sup>o</sup> | t = 3.72 | 18 | 0.005* |  |  |

\* Statistically significant value.
<sup>ⓐ</sup> Sidak post hoc comparison, alfa adjusted by Bonferroni at 0.05 significance level.

**Supp. Table 3.** Statistical data corresponding to Figure 3, including the statistical test, sample size (n), value of the statistic, degrees of freedom (df), associated *p* value, effect size (η _p_ ^2^) and power effect (1-β).

| Panel | Analysis/group | Statistic | Value | df | p value | $\eta_p^2$ | 1-β |
| --- | --- | --- | --- | --- | --- | --- | --- |
| d | Between groups, PreCNO | One way ANOVA | F = 2.9 | 2,15 | 0.08 | - | - |
|  | Between groups, PostCNO | Two ways ANOVA, RM | F = 116.4 | 2,15 | <0.0001* | 0.93 | 1 |
|  | Sham vs PV-Gi | *post hoc*<sup>Ⓢ</sup> | t = 8.96 | 30 | <0.0001* |  |  |
|  | PV-Gi vs PV-Gq | *post hoc*<sup>Ⓢ</sup> | t = 10.85 | 30 | <0.0001* |  |  |
|  | Sham vs PV-Gq | *post hoc*<sup>Ⓢ</sup> | t = 1.89 | 30 | 0.48 |  |  |
|  | Within groups, Pre-PostCNO | Two ways ANOVA, RM | F = 93.69 | 1,15 | <0.0001* | 0.86 | 1 |
|  | Sham (n=6) | *post hoc*<sup>Ⓢ</sup> | t = 2.18 | 15 | 0.13 |  |  |
|  | PV-Gi (n=6) | *post hoc*<sup>Ⓢ</sup> | t = 17.67 | 15 | <0.0001* |  |  |
|  | PV-Gq (n=6) | *post hoc*<sup>Ⓢ</sup> | t = 3.09 | 15 | 0.02* |  |  |
| e | Between groups, PreCNO | One way ANOVA | F = 0.45 | 2,15 | 0.64 | - | - |
|  | Between groups, PostCNO | Two ways ANOVA, RM | F = 68.12 | 2,15 | <0.0001* | 0.9 | 1 |
|  | Sham vs PV-Gi | *post hoc*<sup>Ⓢ</sup> | t = 14.08 | 30 | <0.0001* |  |  |
|  | PV-Gi vs PV-Gq | *post hoc*<sup>Ⓢ</sup> | t = 15.16 | 30 | <0.0001* |  |  |
|  | Sham vs PV-Gq | *post hoc*<sup>Ⓢ</sup> | t = 1.07 | 30 | 0.64 |  |  |
|  | Within groups, Pre-PostCNO | Two ways ANOVA, RM | F = 63.82 | 1,15 | <0.0001* | 0.81 | 1 |
|  | Sham (n=6) | *post hoc*<sup>Ⓢ</sup> | t = 0.28 | 15 | 0.98 |  |  |
|  | PV-Gi (n=6) | *post hoc*<sup>Ⓢ</sup> | t = 14.13 | 15 | <0.0001* |  |  |
|  | PV-Gq (n=6) | *post hoc*<sup>Ⓢ</sup> | t = 0.57 | 15 | 0.92 |  |  |
| f | Between groups, PostCNO | Two ways ANOVA, RM | F = 0.41 | 2,13 | 0.67 | - | - |
|  | Within groups, Pre-PostCNO | Two ways ANOVA, RM | F = 0.68 | 1,13 | 0.42 | - | - |

\* Statistically significant value.
<sup>Ⓢ</sup> Sidak post hoc comparison, alfa adjusted by Bonferroni at 0.05 significance level.

**Supp. Table 4.** Statistical data corresponding to Figure 4, including the statistical test, sample size (n), value of the statistic, degrees of freedom (df), associated *p* value, effect size (r^2^ or η_p_^2^) and power effect (1-β).

| Panel | Analysis/group | Statistic | Value | df | p value | r<sup>2</sup> | 1-β |
| --- | --- | --- | --- | --- | --- | --- | --- |
| d | Pre-PostCNO, Sham (n=6) | t-test dependent measures | t = 0.88 | 5 | 0.41 | - | - |
|  | Pre-PostCNO, PV-Gi (n=6) |  | t = 4.32 | 5 | 0.0076* | 0.79 | 0.92 |
|  | Pre-PostCNO, PV-Gq (n=5) |  | t = 1.86 | 4 | 0.16 | - | - |
| e | Pre-PostCNO, Sham (n=6) | t-test dependent measures | t = 1.57 | 5 | 0.18 | - | - |
|  | Pre-PostCNO, PV-Gi (n=6) |  | t = 8.12 | 5 | 0.0005* | 0.93 | 0.99 |
|  | Pre-PostCNO, PV-Gq (n=5) |  | t = 0.82 | 4 | 0.45 | - | - |
| f | Pre-PostCNO, PV-Gi (n=6) | t-test dependent measures | t = 3.43 | 5 | 0.01* | 0.7 | 0.78 |
| g | Pre-PostCNO, PV-Gi (n=6) | t-test dependent measures | t = 2.17 | 5 | 0.08 | 0.48 | 0.42 |
| $\eta_p^2$ | | | | | | | |
| h | Pre-PostCNO, Sham (n=6) | Two ways ANOVA, RM | F = 1.57 | 1,50 | 0.21 | - | - |
|  | Pre-PostCNO, PV-Gi (n=4)<sup>ψ</sup> |  | F = 86.73 | 1,30 | <0.0001* | 0.74 | 0.98 |
|  | Pre-PostCNO, PV-Gq (n=6) |  | F = 0.7 | 1,50 | 0.4 | - | - |
| j | Between groups. All areas.<br>Sham (n=8)<br>PV-Gi (n=9)<br>PV-Gq (n=6) | Two ways ANOVA | F = 0.21 | 2,91 | 0.8 | - | - |

\* Statistically significant value.
<sup>ψ</sup> Two animals were excluded due to incorrect anatomical site of recording in Prefrontal Cortex.

**Supp. Table 5.**
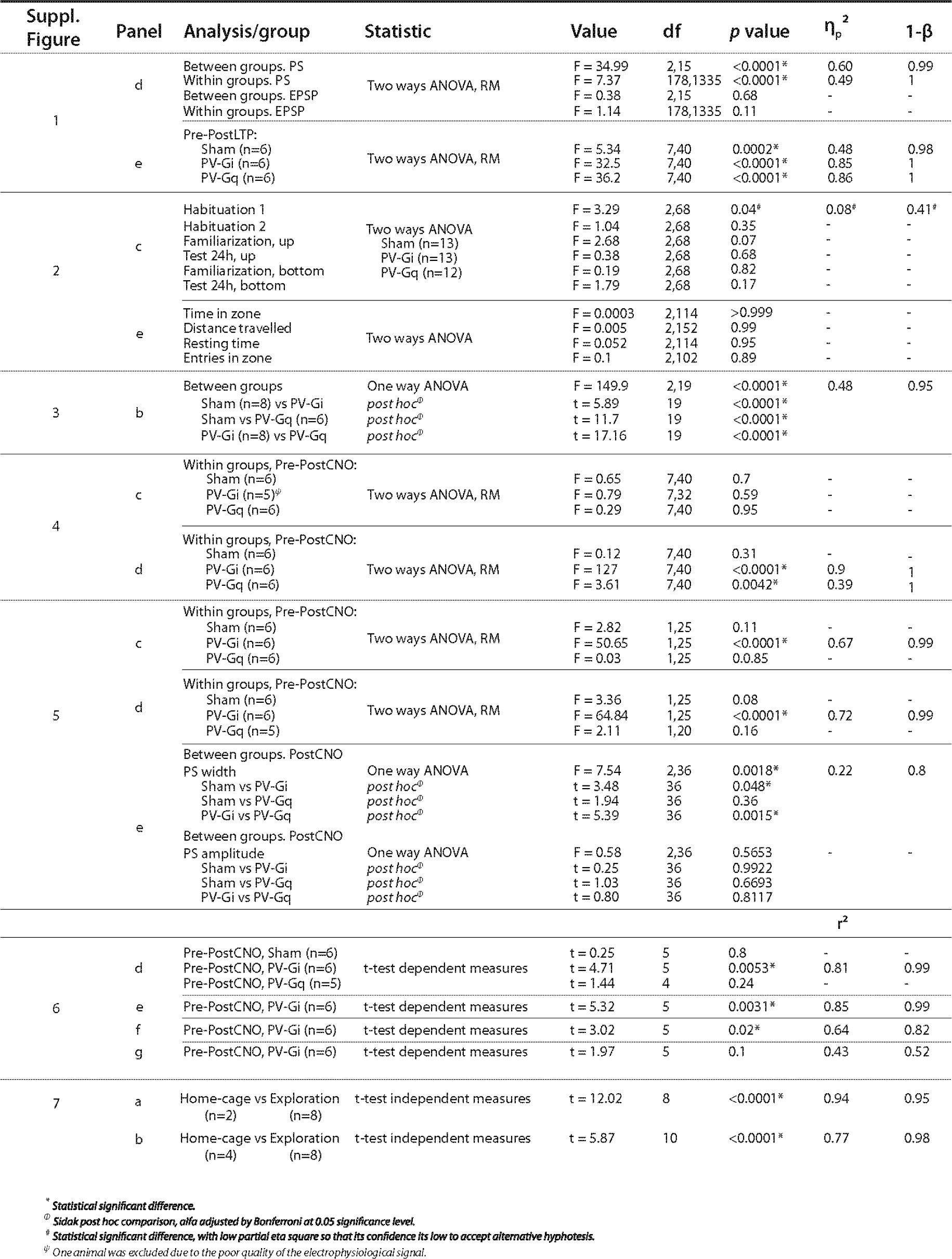
Statistical data corresponding to Supp. Figures 1-7, including the statistical test, sample size (n), value of the statistic, degrees of freedom (df), associated *p* value, effect size (η _p_ ^2^ or r^2^) and power effect (1-β).

## Notes

### Competing Interest Statement

The authors have declared no competing interest.

## References

1. S. Tonegawa, M. D. Morrissey, T. Kitamura, The role of engram cells in the systems consolidation of memory. Nat. Rev. Neurosci. 19, 485–498 (2018).

2. D. Marr, Simple memory: a theory for archicortex. Philos. Trans. R. Soc. Lond. B. Biol. Sci. 262, 23–81 (1971).

3. A. Treves, E. T. Rolls, Computational constraints suggest the need for two distinct input systems to the hippocampal CA3 network. Hippocampus. 2, 189–99 (1992).

4. J. K. Leutgeb, S. Leutgeb, M.-B. Moser, E. I. Moser, Pattern Separation in the Dentate Gyrus and CA3 of the Hippocampus. Science (80-.). 315, 961–966 (2007).

5. S. Ramirez, X. Liu, P.-A. Lin, J. Suh, M. Pignatelli, R. L. Redondo, T. J. Ryan, S. Tonegawa, Creating a False Memory in the Hippocampus. Science (80-.). 341, 387–391 (2013).

6. S. Daumas, H. Halley, B. Francés, J. M. Lassalle, Encoding, consolidation, and retrieval of contextual memory: Differential involvement of dorsal CA3 and CA1 hippocampal subregions. Learn. Mem. 12, 375–382 (2005).

7. F. Xia, B. A. Richards, M. M. Tran, S. A. Josselyn, K. Takehara-Nishiuchi, P. W. Frankland, Parvalbumin-positive interneurons mediate neocortical-hippocampal interactions that are necessary for memory consolidation. Elife. 6 (2017), doi: 10.7554/eLife.27868.

8. L. R. Squire, P. Alvarez, Retrograde amnesia and memory consolidation: a neurobiological perspective. Curr. Opin. Neurobiol. 5, 169–177 (1995).

9. P. W. Frankland, B. Bontempi, The organization of recent and remote memories. Nat. Rev. Neurosci. 6, 119–130 (2005).

10. S.-H. Wang, R. G. M. Morris, Hippocampal-Neocortical Interactions in Memory Formation, Consolidation, and Reconsolidation. Annu. Rev. Psychol. 61, 49–79 (2010).

11. S. Canals, M. Beyerlein, H. Merkle, N. K. Logothetis, Functional MRI Evidence for LTP-Induced Neural Network Reorganization. Curr. Biol. 19, 398–403 (2009).

12. E. Alvarez-Salvado, V. Pallares, A. Moreno, S. Canals, Functional MRI of long-term potentiation: imaging network plasticity. Philos. Trans. R. Soc. B Biol. Sci. 369, 20130152–20130152 (2013).

13. G. Del Ferraro, A. Moreno, B. Min, F. Morone, Ú. Pérez-Ramírez, L. Pérez-Cervera, L. C. Parra, A. Holodny, S. Canals, H. A. Makse, Publisher Correction: Finding influential nodes for integration in brain networks using optimal percolation theory. Nat. Commun. 9, 3156 (2018).

14. T. V. P. Bliss, G. L. Collingridge, A synaptic model of memory: Long-term potentiation in the hippocampus. Nature. 361 (1993), pp. 31–39.

15. R. C. Malenka, M. F. Bear, LTP and LTD: An embarrassment of riches. Neuron. 44 (2004), pp. 5–21.

16. G. Neves, S. F. Cooke, T. V. P. Bliss, Synaptic plasticity, memory and the hippocampus: A neural network approach to causality. Nat. Rev. Neurosci. 9 (2008), pp. 65–75.

17. R. A. Nicoll, R. C. Malenka, Contrasting properties of two forms of long-term potentiation in the hippocampus. Nature. 377, 115–118 (1995).

18. S. J. Martin, P. D. Grimwood, R. G. M. Morris, Synaptic Plasticity and Memory: An Evaluation of the Hypothesis. Annu. Rev. Neurosci. 23, 649–711 (2000).

19. J. Z. Young, A model of the brain (Oxford, 1964).

20. N. McNaughton, R. G. M. Morris, Chlordiazepoxide, an anxiolytic benzodiazepine, impairs place navigation in rats. Behav. Brain Res. 24, 39–46 (1987).

21. M. P. Arolfo, J. D. Brioni, Diazepam impairs place learning in the Morris water maze. Behav. Neural Biol. 55, 131–136 (1991).

22. I. Izquierdo, J. H. Medina, M. Bianchin, R. Walz, M. S. Zanatta, R. C. Da Silva, M. B. E. Silva, A. C. Ruschel, N. Paczko, Memory processing by the limbic system: Role of specific neurotransmitter systems. Behav. Brain Res. 58, 91–98 (1993).

23. N. Collinson, J. R. Atack, P. Laughton, G. R. Dawson, D. N. Stephens, An inverse agonist selective for α5 subunit-containing GABAA receptors improves encoding and recall but not consolidation in the Morris water maze. Psychopharmacology (Berl). 188, 619–628 (2006).

24. M. S. Chambers, J. R. Atack, H. B. Broughton, N. Collinson, S. Cook, G. R. Dawson, S. C. Hobbs, G. Marshall, K. A. Maubach, G. V. Pillai, A. J. Reeve, A. M. MacLeod, Identification of a novel, selective GABAa α5 receptor inverse agonist which enhances cognition. J. Med. Chem. 46, 2227–2240 (2003).

25. J. J. Letzkus, S. B. E. Wolff, A. Lüthi, Disinhibition, a Circuit Mechanism for Associative Learning and Memory. Neuron. 88, 264–276 (2015).

26. R. C. Froemke, Plasticity of Cortical Excitatory-Inhibitory Balance. Annu. Rev. Neurosci. 38, 195–219 (2015).

27. F. Donato, S. B. Rompani, P. Caroni, Parvalbumin-expressing basket-cell network plasticity induced by experience regulates adult learning. Nature. 504, 272–276 (2013).

28. G. M. Alexander, S. C. Rogan, A. I. Abbas, B. N. Armbruster, Y. Pei, J. A. Allen, R. J. Nonneman, J. Hartmann, S. S. Moy, M. A. Nicolelis, J. O. McNamara, B. L. Roth, Remote control of neuronal activity in transgenic mice expressing evolved G protein-coupled receptors. Neuron. 63, 27–39 (2009).

29. P. E. Gilbert, R. P. Kesner, I. Lee, Dissociating hippocampal subregions: double dissociation between dentate gyrus and CA1. Hippocampus. 11, 626–36 (2001).

30. M. Leger, A. Quiedeville, V. Bouet, B. Haelewyn, M. Boulouard, P. Schumann-Bard, T. Freret, Object recognition test in mice. Nat. Protoc. 8, 2531–7 (2013).

31. T. Stefanelli, C. Bertollini, C. Lüscher, D. Muller, P. Mendez, Hippocampal Somatostatin Interneurons Control the Size of Neuronal Memory Ensembles. Neuron. 89, 1074–85 (2016).

32. O. Herreras, J. M. Solis, R. Martin del Rio, J. Lerma, Characteristics of CA1 activation through the hippocampal trisynaptic pathway in the unanaesthetized rat. Brain Res. 413, 75–86 (1987).

33. P. Jego, J. Pacheco-Torres, A. Araque, S. Canals, Functional MRI in mice lacking IP3-dependent calcium signaling in astrocytes. J. Cereb. Blood Flow Metab. 34, 1599–603 (2014).

34. D. Tse, R. F. Langston, M. Kakeyama, I. Bethus, P. A. Spooner, E. R. Wood, M. P. Witter, R. G. M. Morris, Schemas and memory consolidation. Science (80-.). 316, 76–82 (2007).

35. P. A. Vieira, E. Korzus, CBP-Dependent memory consolidation in the prefrontal cortex supports object-location learning. Hippocampus. 25, 1532–1540 (2015).

36. G. R. I. Barker, E. C. Warburton, NMDA Receptor Plasticity in the Perirhinal and Prefrontal Cortices Is Crucial for the Acquisition of Long-Term Object-in-Place Associative Memory. J. Neurosci. 28, 2837–2844 (2008).

37. A. J. D. Nelson, K. E. Thur, C. A. Marsden, H. J. Cassaday, Dissociable Roles of Dopamine Within the Core and Medial Shell of the Nucleus Accumbens in Memory for Objects and Place. Behav. Neurosci. 124, 789–799 (2010).

38. T. Takeuchi, A. J. Duszkiewicz, A. Sonneborn, P. A. Spooner, M. Yamasaki, M. Watanabe, C. C. Smith, G. Fernández, K. Deisseroth, R. W. Greene, R. G. M. Morris, Locus coeruleus and dopaminergic consolidation of everyday memory. Nature. 537, 357–362 (2016).

39. D. E. Berman, Y. Dudai, Memory extinction, learning anew, and learning the new: dissociations in the molecular machinery of learning in cortex. Science. 291, 2417–9 (2001).

40. S. R. Cobb, E. H. Buhl, K. Halasy, O. Paulsen, P. Somogyi, Synchronization of neuronal activity in hippocampus by individual GABAergic interneurons. Nature. 378, 75–78 (1995).

41. R. Miles, K. Tóth, A. I. Gulyás, N. Hájos, T. F. Freund, Differences between somatic and dendritic inhibition in the hippocampus. Neuron. 16, 815–823 (1996).

42. J. Basu, K. V Srinivas, S. K. Cheung, H. Taniguchi, Z. J. Huang, S. A. Siegelbaum, A cortico-hippocampal learning rule shapes inhibitory microcircuit activity to enhance hippocampal information flow. Neuron. 79, 1208–21 (2013).

43. H. Takahashi, J. C. Magee, Pathway Interactions and Synaptic Plasticity in the Dendritic Tuft Regions of CA1 Pyramidal Neurons. Neuron. 62, 102–111 (2009).

44. A. Moreno, R. G. M. Morris, S. Canals, Frequency-Dependent Gating of Hippocampal–Neocortical Interactions. Cereb. Cortex. 26, 2105–2114 (2016).

45. V. S. Sohal, F. Zhang, O. Yizhar, K. Deisseroth, Parvalbumin neurons and gamma rhythms enhance cortical circuit performance. Nature. 459, 698–702 (2009).

46. J. A. Cardin, M. Carlén, K. Meletis, U. Knoblich, F. Zhang, K. Deisseroth, L.-H. Tsai, C. I. Moore, Driving fast-spiking cells induces gamma rhythm and controls sensory responses. Nature. 459, 663–667 (2009).

47. G. Orgy Buzsáki, X.-J. Wang, Mechanisms of Gamma Oscillations. Annu. Rev. Neurosci. 35, 203–25 (2012).

48. P. Fries, Neuronal Gamma-Band Synchronization as a Fundamental Process in Cortical Computation. Annu. Rev. Neurosci. 32, 209–224 (2009).

49. J. Courtin, N. Karalis, C. Gonzalez-Campo, H. Wurtz, C. Herry, Persistence of amygdala gamma oscillations during extinction learning predicts spontaneous fear recovery. Neurobiol. Learn. Mem. 113, 82–89 (2014).

50. V. J. Lopez-Madrona, E. Alvarez-Salvado, D. Moratal, O. Herreras, E. Pereda, C. R. Mirasso, S. Canals, Gamma oscillations coordinate different theta rhythms in the hippocampus. bioRxiv, 418434 (2018).

51. A. J. Rashid, C. Yan, V. Mercaldo, H.-L. L. Hsiang, S. Park, C. J. Cole, A. De Cristofaro, J. Yu, C. Ramakrishnan, S. Y. Lee, K. Deisseroth, P. W. Frankland, S. A. Josselyn, Competition between engrams influences fear memory formation and recall. Science. 353, 383–7 (2016).

52. E. Lesburguères, O. L. Gobbo, S. Alaux-Cantin, A. Hambucken, P. Trifilieff, B. Bontempi, Early tagging of cortical networks is required for the formation of enduring associative memory. Science. 331, 924–8 (2011).

53. T. Kitamura, S. K. Ogawa, D. S. Roy, T. Okuyama, M. D. Morrissey, L. M. Smith, R. L. Redondo, S. Tonegawa, Engrams and circuits crucial for systems consolidation of a memory. Science. 356, 73–78 (2017).

54. J. W. Lee, M. W. Jung, Separation or binding? Role of the dentate gyrus in hippocampal mnemonic processing. Neurosci. Biobehav. Rev. 75, 183–194 (2017).

55. S. R. Tsur, Y. E. Demiray, K. Tripathi, O. Stork, G. Richter-Levin, A. Albrecht, Region-specific involvement of interneuron subpopulations in trauma-related pathology and resilience. Neurobiol. Dis. 143, 104974 (2020).

56. K.B.J Franklin, G. Paxinos, The mouse brain in stereotaxic coordinates (Elsevier Acaemic Press, San Diego, ed. 3rd, 2007).

57. F. de Chaumont, S. Dallongeville, N. Chenouard, N. Hervé, S. Pop, T. Provoost, V. Meas-Yedid., P. P., T. Lecomte, Y. Le Montagner, T. Lagache, A. Dufour, D. C. Olivo-Marin, Icy: an open bioimage informatics platform for extended reproducible research. Nat. Methods. 9, 690–696 (2012).

58. R. A. Fisher, Statistical Methods for Research Workers (Oliver and Boyd, Edinburgh, Scotland, 1925).

